# Non-living respiration: another breath in the soil?

**DOI:** 10.1101/2025.07.03.662961

**Authors:** Clémentin Bouquet, Benoit Kéraval, Mounir Traikia, Gael Alvarez, Fanny Perrière, Anne-Hélène Le Jeune, Hermine Billard, Jonathan Colombet, Sandrine Revaillot, Sébastien Fontaine, Anne-Catherine Lehours

**Affiliations:** Université Clermont Auvergne, CNRS, LMGE, F-63000 Clermont-Ferrand, France; Université Clermont Auvergne, CNRS, ICCF, F-63000 Clermont-Ferrand, France; Université Clermont Auvergne, PFEM, MetaboHUB Clermont, Clermont-Ferrand, France; Université Clermont Auvergne, INRAE, VetAgro Sup, UMR Ecosystème Prairial, 63000 Clermont-Ferrand, France

## Abstract

The present study challenges the traditional view that respiration of organic carbon to CO_2_ is exclusively an intracellular process, revealing that organic compound respiration can occur spontaneously in an extracellular context in soils.

Using ^1^H nuclear magnetic resonance spectroscopy to analyse the dynamics of the sterile soil exometabolomes alongside C-CO_2_ flux analyses and sterile soil fuel cells, we show that soil catalysts facilitate a diverse array of substrate-driven reactions, leading to the complete oxidation of organic compounds to CO_2_ with O_2_ consumption.

Our results indicate that soil particles are capable of transferring electrons from substrates to the final acceptor, sustaining metabolic processes independently of living cells. Notably, some soil catalysts and induced respiration remain stable for over six years.

Our results support the coexistence of cellular and non-cellular metabolic pathways in soil respiration.

## INTRODUCTION

In a universe that is constantly tending towards greater disorder, living organisms must actively create and maintain order to survive, a trait that fundamentally distinguishes them from non-living matter. To combat this inherent drive towards entropy, cells must carry out a never-ending stream of chemical reactions. In heterotrophic cells, the essence of these cascades of redox reactions, known as respiration, is the oxidation of small organic molecules (*e*.*g*., organic acids, amino acids, and sugars) that fuel cells with the energy stored in their chemical bonds and provide many other molecules that the cell needs. The captured electrons are transferred to a terminal electron acceptor, namely molecular oxygen (O_2_) in aerobic respiration, and CO_2_ and H_2_O are the end products of these reactions. These cellular biochemical reactions are organised in such a way as to make them possible at low temperatures and to recover the energy released.

In addition to the functions of respiration in cell bioenergetics, this cellular mechanism is crucial for forecasting the global carbon cycle and its climate feedbacks. In terrestrial ecosystems, in particular, soil respiration is the primary process by which CO_2_ fixed by photosynthetic organisms in organic matter (OM) is recycled into the atmosphere (**Raich & Schlesinger 1992; Nissan *et al*. 2023**). With soil OM containing three times as much carbon as the atmosphere, a small change in soil respiration can have a major impact on atmospheric CO_2_ and the earth’s temperature (**Cox *et al*. 2000; Bond-Lamberty & Thomson 2010, Hicks Pries *et al*. 2017**). This has prompted scientists since the early twentieth century to study the catabolic activities of soil microbes and their response to environmental factors (**Lundegårdh 1927**). Surprisingly, several studies have found that soil respiration is not always connected to size and composition of the microbial biomass, as experimental reductions in these microbial components have moderate effects on the respiration rate (*e*.*g*., **Kemmitt *et al*. 2008**). It has also been observed that substantial CO_2_ emissions can persist for several weeks in soils where microbial life has been reduced to an undetectable value by exposure to toxics (ClCH_3_, orange acridine), *gamma irradiation*, or autoclaving (**Ramsay & Bawden 1983; Lensi *et al*. 1991; Trevors 1996; Kemmitt *et al*. 2008; Maire *et al*. 2013; Kéraval *et al*. 2016**). Such CO_2_ emissions cannot be explained by an activity of surviving microbes that would be present in small quantities in soils, unless we consider unrealistic cellular respiratory activity (**Maire *et al*. 2013**). Furthermore, the isotopic signature of CO_2_ emissions (δ^13^C =-75.4 ± 2.8 ‰) from sterilised soils with excess of substrates indicates an isotope fractionation during the conversion process of organic C (δ^13^C = -27.4 ± 0.4 ‰) to CO_2_ (**Kéraval *et al*. 2016**) incompatible with cell-derived respiration (**Santruckova *et al*. 2000; Wang *et al*. 2015**).

Various noncellular processes have been proposed as potential contributors to CO_2_ emissions in sterile soils such as (i) single catalytic reactions induced by reactive oxygen species (ROS) and/or metals such as iron (**Yu & Kuzyakov 2021**) or supported by decarboxylases released during cell lysis (**Maire *et al*. 2013; Blankinship *et al*. 2014**), (ii) complete mineralisation of organic compounds supported by extracellular oxidative metabolisms (EXOMETs; **Maire *et al*. 2013; Kéraval *et al*. 2016**). EXOMETs are distinguished from the other processes by their complexity and persistence over time. EXOMETs, according to the definition proposed by **Maire *et al*. (2013)**, involve numerous coupled redox reactions carried out by soil-stabilised enzymes and/or abiotic catalysts (clay, metals) reconstituting an equivalent of a cell respiration process capable of converting organic compounds such as glucose into CO_2_ with electron transfer to O_2_. The EXOMETs hypothesis is challenging. The respiratory machinery is rather complicated and involves numerous respiratory-mediating endoenzymes whose functioning depends on a variety of redox cofactors that need to be regenerated (*e*.*g*., NAD+), their location close to other enzymes, and the physiological properties of the cell (*e*.*g*., redox potential) (**Efremov & Sazanov 2011)**. Owing to this complexity and the fact that cellular enzymes are superbly crafted but vulnerable catalysts, respiration is currently considered to be a strictly intracellular metabolic process.

In the present study, we have combined exometabolomics and isotope approaches, in medium-term (∼ 6 months) and long-term (> 6 years) experiments in soil microcosms, as well as the construction of fuel cells using sterile soils, to demonstrate that soils contain catalysts that are stable over years and capable of generating a chain of redox reactions that lead to the complete mineralisation of various organic compounds to CO_2_ and electron transfer to oxygen. These results show that the respiration of organic compounds is not a cell-specific process and that it can spontaneously occur in an extracellular context in soils.

## RESULTS and DISCUSSION

### Non-cellular processes generate both the production and consumption of various metabolites

To study whether the sole action of soil catalysers is able to sustain chains of chemical reactions, we examined the exometabolomes of sterilised and non-sterilised soils incubated for 163 days in the dark at 20 °C. The soils were sterilised by *gamma irradiation* at 45 kGy, which is an effective sterilising agent in the soil particularly at doses above 20 kGy (**Denisova *et al*. 2005**), and does not disrupt soil structure and chemistry as much as autoclaving (**McNamara *et al*. 2003, Lees *et al*. 2018**). Three substrate treatments were applied to sterilised soils to test the ability of soil catalysers to degrade different substrates of varying complexity: no substrate addition (IS), the addition of ^13^C-Citrate (C-IS) or ^13^C-Glucose (G-IS) **(Appendix S1: Section S1)**. All manipulations were carried out under strict sterile conditions and the sterility of IS, C-IS, and G-IS samples was checked using live / dead cell staining coupled to flow cytometry analysis **(Appendix S1: Section S2 and Figure S.1)**, an approach previously validated in this research (**Kéraval *et al*. 2016; 2018**). The extracellular fraction of molecules that are inferred to be produced and / or used in soil, i.e. exometabolome, was profiled at six dates during incubation using untargeted metabolomics, a robust approach to provide comprehensive analyses of complex extracts **(Swenson *et al*. 2015, Song *et al*. 2024**). We use proton nuclear magnetic resonance spectroscopy (NMR) to track the dynamics of the water-extractable fraction of exometabolites with a molecular weight < 3 kDa (**Appendix S1: Section S3)**. Water is an excellent solvent for examining total extractable OM in soils (**Swenson *et al*. 2015**) and the 0 to 3 kDa size class of metabolites, including small to mid-sized molecules (*e*.*g*. amino acids, carbohydrates, organic acids), represents most of the compounds of dissolved OM in soils (**Malik & Gleixner 2013**). Chemical information encoded in NMR data was explored after manual grouping (**Beckonert *et al*. 2007, Marchand *et al*. 2018**) of the spectral responses into variable size buckets (*i*.*e*. a spectral region) subjected to statistical data analysis after a normalisation step (**Appendix S1: Section S3)**.

A total of 177 and 506 distinct buckets between 0 and 10 ppm NMR spectra were identified in the exometabolome of non-sterilised (LS) and IS samples, respectively (**Appendix S1: Figure S2)**. At the start of incubation (T1), the molecular richness and diversity were significantly higher in IS samples than in LS samples (**Figs.1A and 1B**), presumably because cellular metabolites were released from dead biomass after *gamma irradiation*. In LS samples, both indicators decreased during incubation (**Figs.1A and 1B**), consistent with evidence that microbial decomposition reduces the molecular diversity of soil OM (**Davenport *et al*. 2023**).

**Figure 1:**
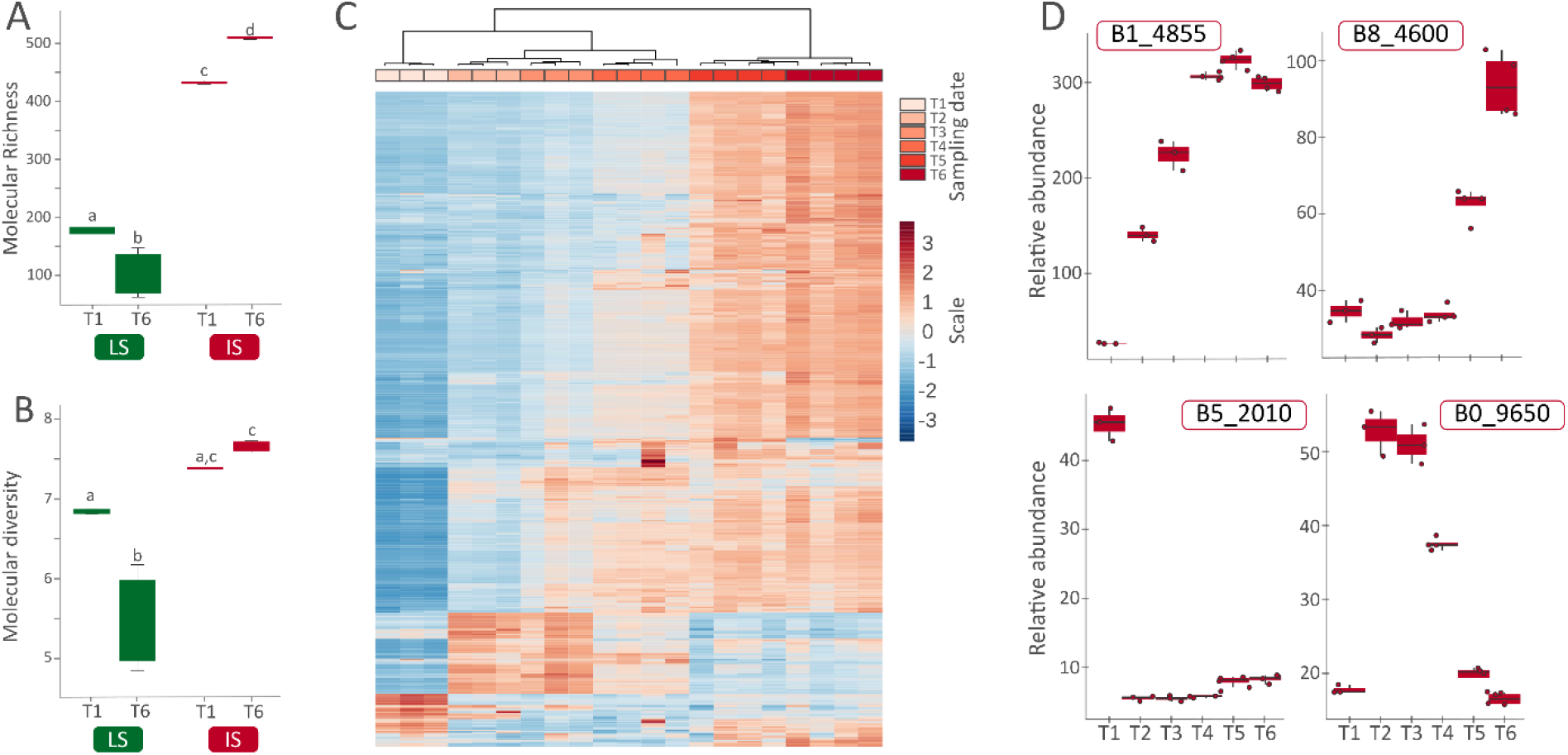
Richness and diversity of the global exometabolome in non-sterilised (LS) and sterilised (IS) soils and dynamics of selected buckets in IS microcosms. **(A)** Molecular richness (number of non-null buckets) and **(B)** molecular diversity (expressed by the Shannon diversity index) of the water-extractable fraction of exometabolites with molecular weight < 3 kDa in non-sterilised soil (LS) and sterilised soil (IS) at the beginning (T1 = 0.2 day) and end (T6 = 163 days) of the incubation period. To test for differences between two groups, the mixed effects model was applied using the Holm-Sidak post hoc test. **(C)** Heat map showing the relative intensities of the water-extractable fraction of exometabolites with molecular weights < 3 kDa in sterilised soil (IS) at the different sampling dates. The relative abundance is normalised to the concentration of the reference molecule during ^1^H-NMR processing. The shades of red and blue indicate increasing and decreasing intensities, respectively. The colour scale represents the magnitude of the change. Individual samples are plotted on the horizontal axis, and buckets are plotted on the vertical axis. The Euclidean distance metric and Ward’s clustering method were used for hierarchical clustering of the samples. **(D)** Temporal dynamics of the relative abundance of four selected buckets (B1-4855; B8_4600; B5-2010, B0-9650) in sterilised soil (IS). Relative abundance is normalised to the concentration of the reference molecule during ^1^H-NMR processing. The name of the bucket corresponds to its mean chemical shift (for example, B0_9560 refers to the bucket identified with a mean chemical shift of 0.9560 ppm). Data are presented as the mean, minimum, and maximum for the four replicate samples by sampling date. T1= 0.2 day, T2=3 days, T3=6 days, T4=17 days, T5=100 days, T6=163 days.

In contrast, exometabolite diversification was promoted over time in sterilised soil samples (**Figs.1A to 1B**), demonstrating that the processes governing metabolite dynamics in the LS and IS microcosms are quite different. Although a general trend towards exometabolite accumulation was observed in the IS, C-IS, and G-IS microcosms between T1 and T6 (**Fig.1C**, **Appendix S1: Figure S3**), bucket dynamics also showed a decrease in some exometabolites over time in three sterilised treatments (**Fig.1C**, **Appendix S1: Figure S3**). Interestingly, some buckets show changing dynamics over time, increasing and then decreasing or vice versa, suggesting that molecules released by some processes are taken up by others (**Fig.1D**). Sorption or desorption of molecules from the solid phase of the soil could contribute to the short-term dynamics, but not over a period of 160 days. Indeed, studies of sorption-desorption kinetics for a wide range of chemical compounds show a stabilisation of the processes within a few hours-days **(Swenson *et al*. 2015; Kleber *et al*. 2021; Spohn *et al*. 2022**). We conclude that the catalysts present in sterilised soils are capable of sustaining a wide range of chemical reactions that produce and consume molecules over several months. Some of these reactions can involve the exchange of molecules, creating metabolic chains in an extracellular context.

### Chemical reactions in sterilised soils have ordered temporal dynamics controlled by substrates

The temporal dynamics of exometabolome fingerprints in sterilised soils (IS, C-IS, and G-IS) and non-sterilised soils (LS), were represented using unsupervised two-dimensional principal component analysis (PCA), with each point in the PCA score plots representing an individual replicate sample (**Fig.2**). Irrespective of sampling date, LS samples showed different exometabolome profiles compared to sterilised soils (**Fig.2A**). For IS, C-IS, and G-IS, a high degree of exometabolome consistency was observed between independent replicates at each sampling date (**Fig.2B; Appendix S1: Figure S4**). In contrast, in LS, biological variation resulted in greater heterogeneity between biological replicates, particularly in the later stages of incubation (**Appendix S1: Figure S5**).

**Figure 2:**
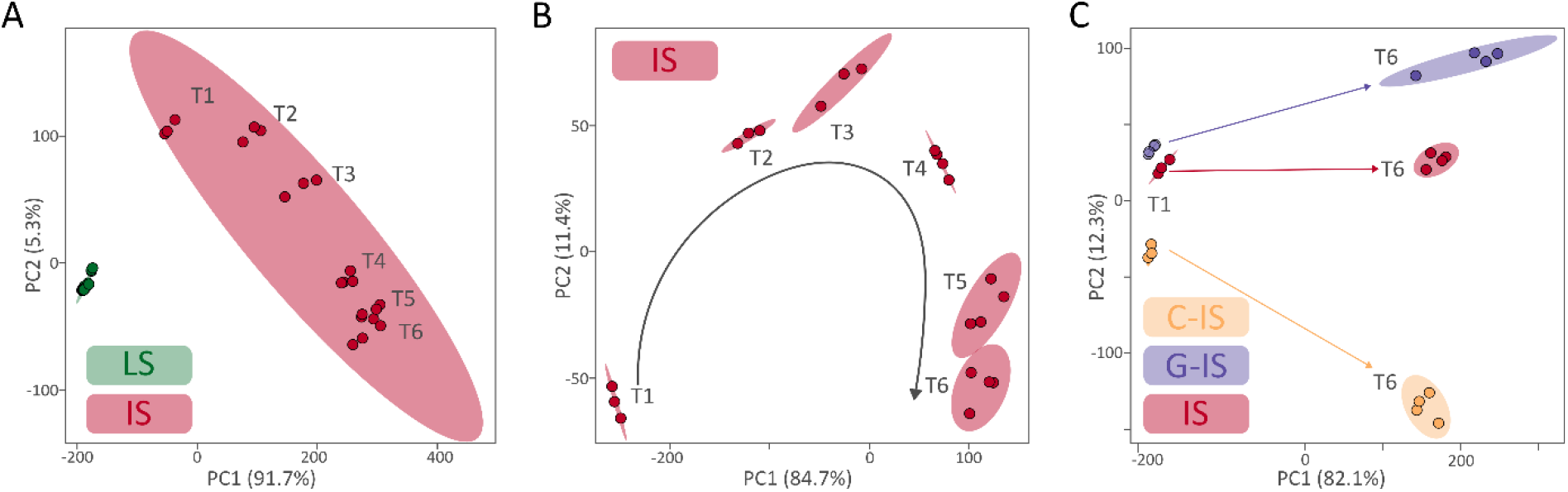
Dynamics of the water-extractable fraction of exometabolites with molecular weight < 3 kDa in soil microcosms, analysed by proton nuclear magnetic resonance spectroscopy (^1^H-NMR) **(A)** Principal component analysis (PCA) of the temporal dynamics of the exometabolome in non-sterilised soils (LS) and irradiated soils (IS) between T1 and T6. For LS, all dates and replicates are represented but overlap. **(B)** PCA of the temporal dynamics of the exometabolome in IS samples between T1 and T6. **(C)** PCA of the dynamics of the exometabolome at T1 (0.2 days) and T6 (163 days) for IS, and for sterilised soils amended with ^13^C-Glucose (G-IS) or ^13^C-Citrate (C-IS). To make the graph easier to read, only the dates T1 and T6 are shown, but the PCA plots for C-IS and G-IS are shown in supplementary material S.4. Each point on the PCA corresponds to a sacrificed replicate (4 replicates per date and per condition) incubated in the dark at 20°C. Coloured ellipses indicate 95 % confidence regions. The two-dimensional PCA score plot shows that the first two principal components account for > 94 % of the variance explained with the first principal component (PC1) retaining > 82.1% of the variance. The arrows in plots B and C have been manually added to highlight the temporal dynamics. T1= 0.2 day, T2=3 days, T3=6 days, T4=17 days, T5=100 days, T6=163 days.

In sterilised soils, the NMR data show time-ordered trajectories of metabolic fingerprint dynamics that diverge progressively between T1 and T6, with the exometabolome composition at one sampling date being closer to that of the previous and subsequent dates than to the others (**Fig.2B; Appendix S1: Figure S4**). Chemistry offers hundreds of different possibilities; such reaction sequences are far from trivial, not least because reaction networks operating under time control are one of the fundamental principles of the functioning of living systems (**Dhasaiyan *et al*. 2022**). The addition of ^13^C-glucose and ^13^C-citrate (3 gC. Kg^-1^ soil), representing less than 10 % of the soil C content (39 gC. Kg^-1^ soil, **Appendix S1: Section S1**), resulted in a significant divergence of exometabolite fingerprints compared to unamended sterilised soil **(Fig. 2C**). Furthermore, exometabolomes and their trajectories diverged between G-IS and C-IS. These substrate-specific responses of exometabolomes indicate that the chemical reactions chains present in sterilised soils are, at least for some of them, capable of processing and transforming different carbon compounds.

### Metabolic activities in sterilised soils lead to complete oxidation of the carbon substrate and consumption of oxygen

Soil catabolic activities were also quantified for non-sterilised soils (LS) and sterilised soils without (IS) and with ^13^C-citrate (C-IS) and ^13^C-glucose (G-IS) by measuring the CO_2_ and O_2_ concentrations, as well as the isotopic signature of C-CO_2_ (δ^13^C-CO_2_) in incubated flasks throughout the 163 days of incubation (**Appendix S1: Section S4**). Not surprisingly, non-sterilised soils induced CO_2_ production and O_2_ consumption (data now shown). Interestingly, sterilised soils also induced both gas exchanges (**Fig. 3**) while the addition of ^13^C-labelled citrate or glucose induced a large increase in δ^13^C-CO_2_ values (**Figs.3A and 3B)** compared to unamended soils (**Fig. 3C**). The δ^13^C-CO_2_ values increased from 330 ± 0.8 to 685.3 ± 0.8 ‰ and from 84.0 ± 0.8 to 182.7 ± 0.9 ‰ between T1 and T6 under glucose and citrate treatments, respectively (**Figs.3A and 3B**), suggesting that glucose and citrate are not directly decarboxylated in sterilised soils but undergo several transformations before being released as CO_2_. These results indicate that the chains of chemical reactions (**Figs. 1 and 2)** carried out by catalysts in sterilised soils can lead to the complete oxidation of molecules such as citrate, glucose, and various soil compounds.

**Figure 3:**
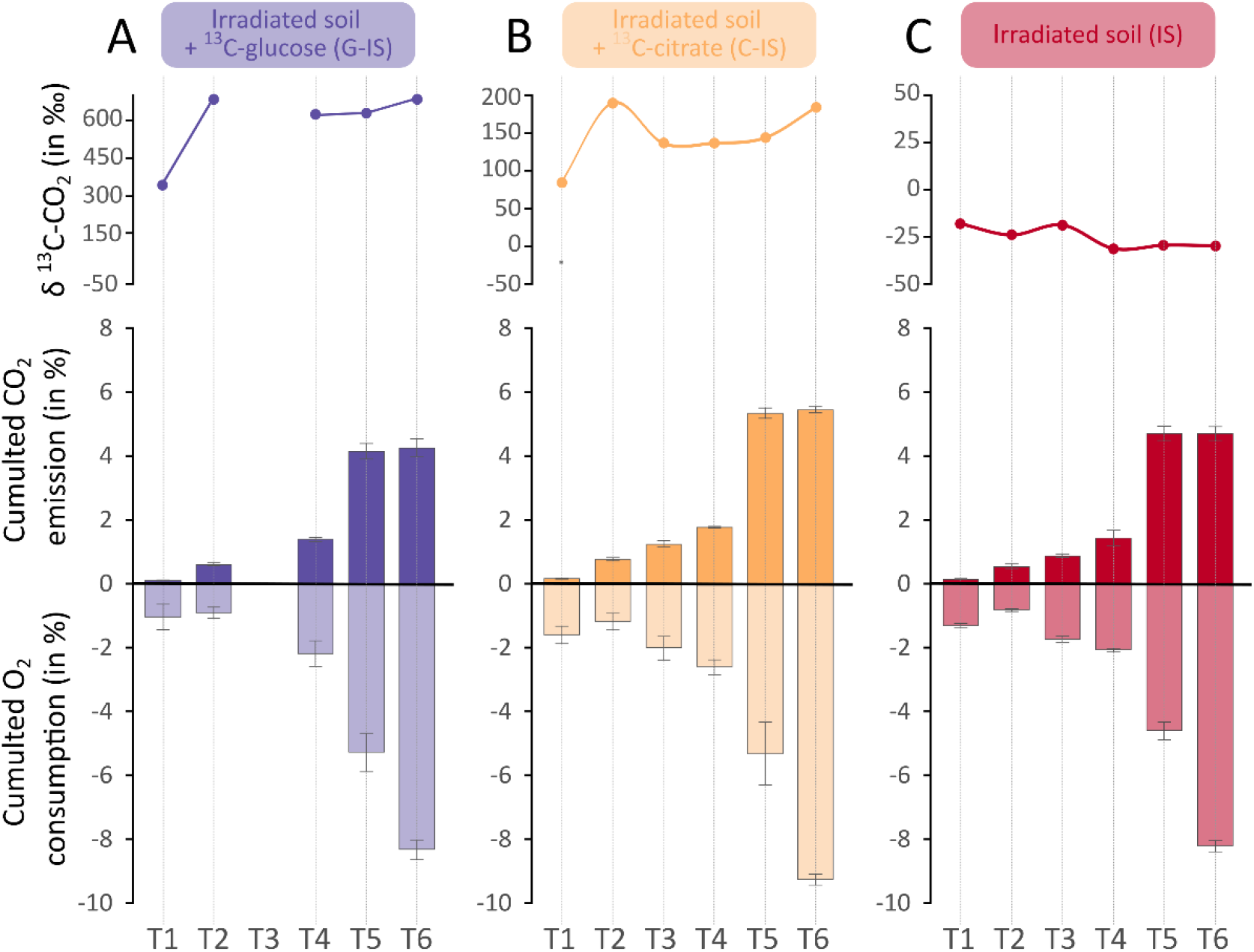
Isotopic signatures of labelled and unlabelled CO_2_, CO_2_ emission, and O_2_ consumption from sterilised soil microcosms amended with ^13^C-glucose (G-IS), ^13^C-citrate (C-IS) or unamended (IS). The isotopic signature (δ^13^C-CO_2_) of CO_2_ is shown at the top of the graph in panels **(A), (B)** and **(C)** with **(A)** irradiated soil amended with ^13^C-glucose (G-IS), **(B)** irradiated soil amended with ^13^C-citrate (C-IS) and **(C)** irradiated unamended soil (IS). Results are expressed in ‰. Total CO_2_ emissions and O_2_ consumption are shown in the lower graphs **(A), (B)** and **(C)**. Results are expressed as a percentage of the microcosm atmosphere. For each sampling date (T1, T2, T3, T4, T5 and T6) and for each parameter, the mean of the values from four replicate samples and the standard deviation are shown. Information on the T3 date for G-IS was not measured. Standard deviations are shown in all graphs. T1= 0.2 day, T2=3 days, T3=6 days, T4=17 days, T5=100 days, T6=163 days.

We note that CO_2_ emission is accompanied with O_2_ consumption under all incubation conditions in sterilised soil (**Fig.3**). The respiratory quotient, which varied from 0.51 to 1.02 as a function of time and experimental conditions, is in the range of that obtained by **Maire *et al*. (2013**). Direct oxidation of organic substrates by O_2_, which has a triplet ground state, is rare due to the generally high energy barrier for electron transfer from the organic substrate to the oxidant (**Piera & Backvall 2008**). To overcome the unfavourable kinetics associated with direct aerobic oxidation, cells use a complex machinery involving cascades of reactions, electron transfer molecules such as NADH and FADH_2_ and protein chains that minimise the energy barrier. Our metabolomics study (**Figs. 1 and 2**) suggests that the oxidation of organic substrates and electron transfers to oxygen can also occur via a cascade of redox reactions in sterilised soils. However, this cascade of redox reactions requires the capacity of soil particles and solution to transfer electrons from one reaction to the other towards oxygen, a capacity that remains to be demonstrated.

### The electron transfer capacity of sterilised soils

To establish the electron transfer capacity of sterilised soils, we constructed an experimental fuel cell (**Fig.4A**) inspired by soil microbial fuel cells (**Barbato *et al*. 2017**). The principle of our single-chamber fuel cell is based on a column of sterilised soil (**Appendix S1: Section S5**) where the upper part is exposed to oxygen from the air, while the lower part is confined to the outside, creating a decreasing oxygen gradient from the top of the cell (the cathode) to the bottom (the anode). The sterilised soil was stored at 20°C for 4.5 years before use, a period long enough for the cellular debris to become invisible under the electron microscope (**Kéraval *et al*. 2016**), and for enzymes to be inactivated (**Maire *et al*. 2013**).

**Figure 4:**
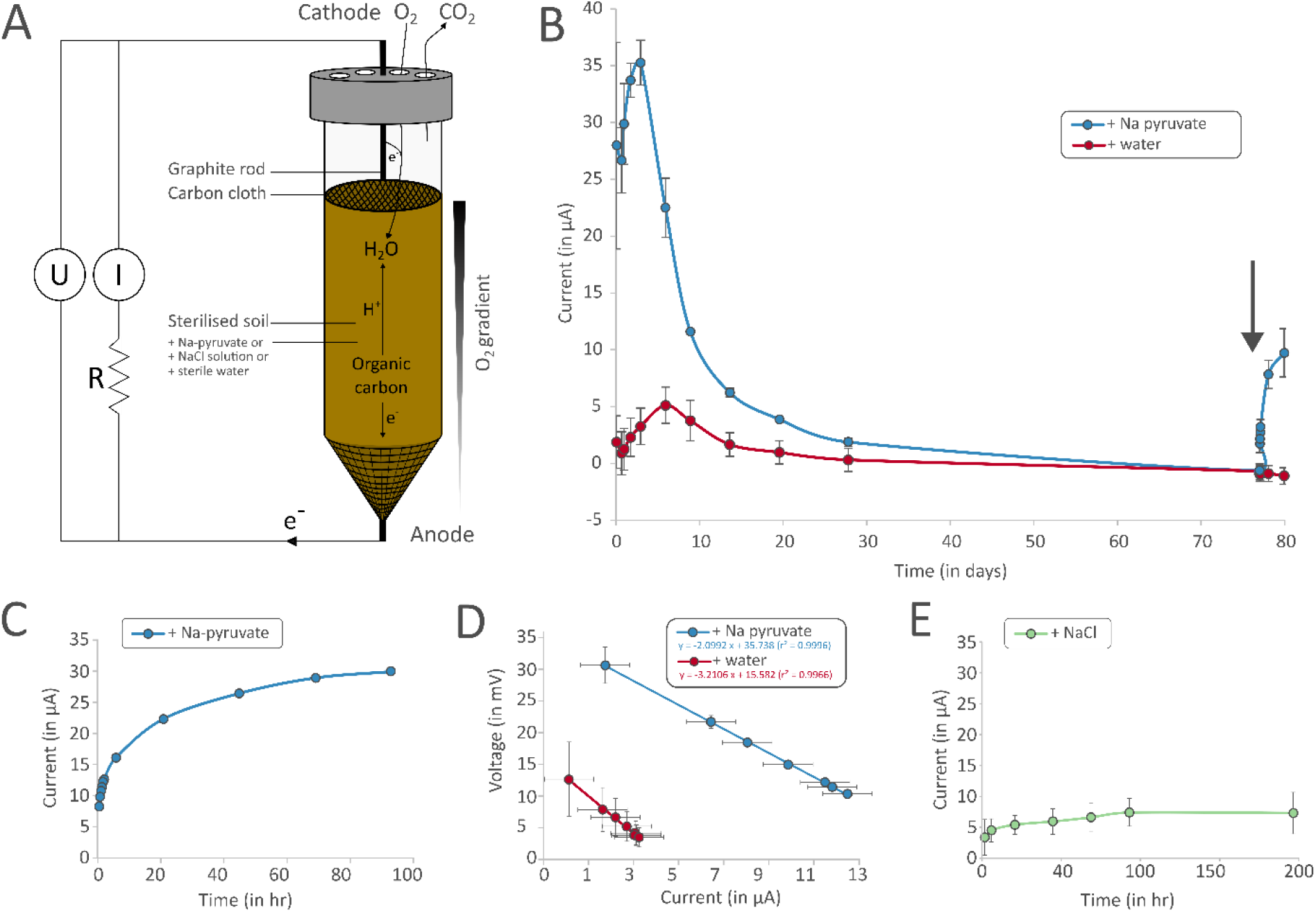
Direct relationship between the electron transfer capacities of sterilised soils and substrate availability revealed by electrical activity measurements of sterilised soil fuel cells. **(A)** Schematic representation of the sterilised soil fuel cell, developed in this study and described in **Supplementary material S.5**. The current induced by the fuel cell was measured after the addition of sterile solutions of sodium pyruvate (Na pyruvate), water, or NaCl at the start of the experiment (T0). The voltage (U) and current (I) generated between the anode and cathode were measured on the external circuit. R and e^-^ represent the circuit resistor and the electrons, respectively. **(B)** Current generated (in µA) as a function of time in sterilised soil fuel cells to which sodium pyruvate (Na pyruvate) or water solutions were added at the start of the experiment (T0) and after 77 days (indicated by the grey arrow). **(C)** Progressive increase in current as a function of time, over the first 96 hours, in a sterilised soil fuel cell to which sodium pyruvate (Na pyruvate) solution was added. **(D)** Cell voltage (in mV) versus resistance (in µA) curves for sterilised soil fuel cells to which sodium pyruvate (Na pyruvate) or water solutions have been added. The electromotive force and resistance of the fuel cell were determined by connecting the fuel cell to decreasing resistances (from 330 to 11900 ohms) at 216 hours. **(E)** Current produced (in µA) as a function of time in sterilised soil fuel cells to which sodium chloride solution was added at the start of the experiment (T0).

When the fuel cell is connected, the oxygen gradient favours the capture of the electrons produced by the oxidation of organic molecules in the buried sterilised soil. The electrons collected at the anode are transported through an external circuit containing a variable resistor to the cathode, where O_2_ acts as an electron acceptor. One substrate treatment (sodium pyruvate, 50 mM) and two control treatments (50 mM sodium chloride or water) were used to verify that the current produced was due to the oxidation of organic molecules. The electricity generated by the fuels cells was characterised by measuring the cell voltage (U) and current (I) across the external circuit by varying the resistance from 330 to 11900 ohms after nine days of fuel cell incubation. The ability of the fuel cells to generate electricity was also regularly checked over a period ranging of 20 minutes to 80 days after construction using a 330 ohm resistance (**Fig. 4B**, **Appendix S1: Section S5**).

Closing the fuel cell circuit induced a significant current in all fuel cells within minutes or hours (**Figs. 4B and 4C**), indicating that soil particles respond rapidly to the oxygen gradient: the soil particles donate electrons to the anode and the soil particles receiving electrons at the cathode ultimately transfer them to the oxygen (**Fig. 4A**). The batteries also show a classic negative linear relationship between voltage and current, which is explained by the internal resistance, given by the slope of the regression line (**Fig. 4D**). Compared to the water treatment, the addition of sodium pyruvate induced a rapid (within 20 minutes) and strong increase in both current and voltage (**Figs. 4B to 4D**). No significant difference in (I) values was found between the two control treatments (water and NaCl, **Appendix S1: Table S1**), confirming sodium pyruvate-induced electricity was promoted by the addition of pyruvate and not by the addition of Na^+^ (**Figs. 4B and 4E**). The current reached a peak (35.3±1.9 µA) three days after the addition of pyruvate and then decreased continuously to reach the control level (**Fig. 4B**). This decrease is related to the depletion of the carbon substrate, as a new addition of pyruvate substrate increased current production (**Fig.4B**). These results show that soil particles mediate electron transfer from the carbon substrate to the final acceptor, in this case, oxygen.

### Long-term dynamics of catabolic activity in sterilised soil are controlled by different catalyst pools

The persistence of non-cellular catabolic activities was studied by incubating microcosms of sterilised soil for 6.7 years. To test whether these activities could be substrate-limited in the long term, we set up two treatments: without and with the addition of ^13^C-glucose (S and S+G respectively) (**Appendix S1: Section S1B**). CO_2_ emission rate and δ^13^C-CO_2_ were measured at nine sampling times over three replicates per condition (**Appendix S1: Section S4**). The sterility of soil microcosms was verified by live / dead staining coupled with flow cytometry analysis, a cultivation approach and observations of thin sections of soil by transmission electron microscopy (**Appendix S1: Section S1)**. At the end of the incubation, no viable cells were detected in (S) and (S+G) (**Appendix S1: Figure S1**).

The kinetics of CO_2_ emission rate in microcosms (S) and (S+G) showed a sharp decrease over the first four days (**Fig. 5A**), from a maximum close to 13 µmol C-CO_2_ day^-1^ after 12 hours of incubation to about 2.4 µmol C-CO_2_ day^-1^ on day 4. The rate of CO_2_ emission then decreased exponentially until it reached 0.27 and 0.34 μmol C-CO_2_ day^-1^ in the (S) and (S+G) microcosms respectively, on day 1125. Between the fourth and sixth year the emission rate tended to stabilise, decreasing by only 4.9 % and 1.5 % per year in microcosms (S) and (S+G), respectively. The addition of ^13^C-glucose in the (S+G) conditions increased the CO_2_ emission rate after four years of incubation compared to the (S) treatment (**Fig. 5A**). This increase in CO_2_ emission rate was associated with the oxidation of ^13^C-glucose, as shown by the incorporation of ^13^C into CO_2_ emissions (**Fig. 5B**). These results indicate that (i) the CO_2_ emission rate of unamended irradiated soil was limited by substrate availability during the latter part of the incubation period and (ii) the ability of soil particles to oxidise glucose and/or its catabolites in an extracellular context can be maintained for more than six years.

**Figure 5:**
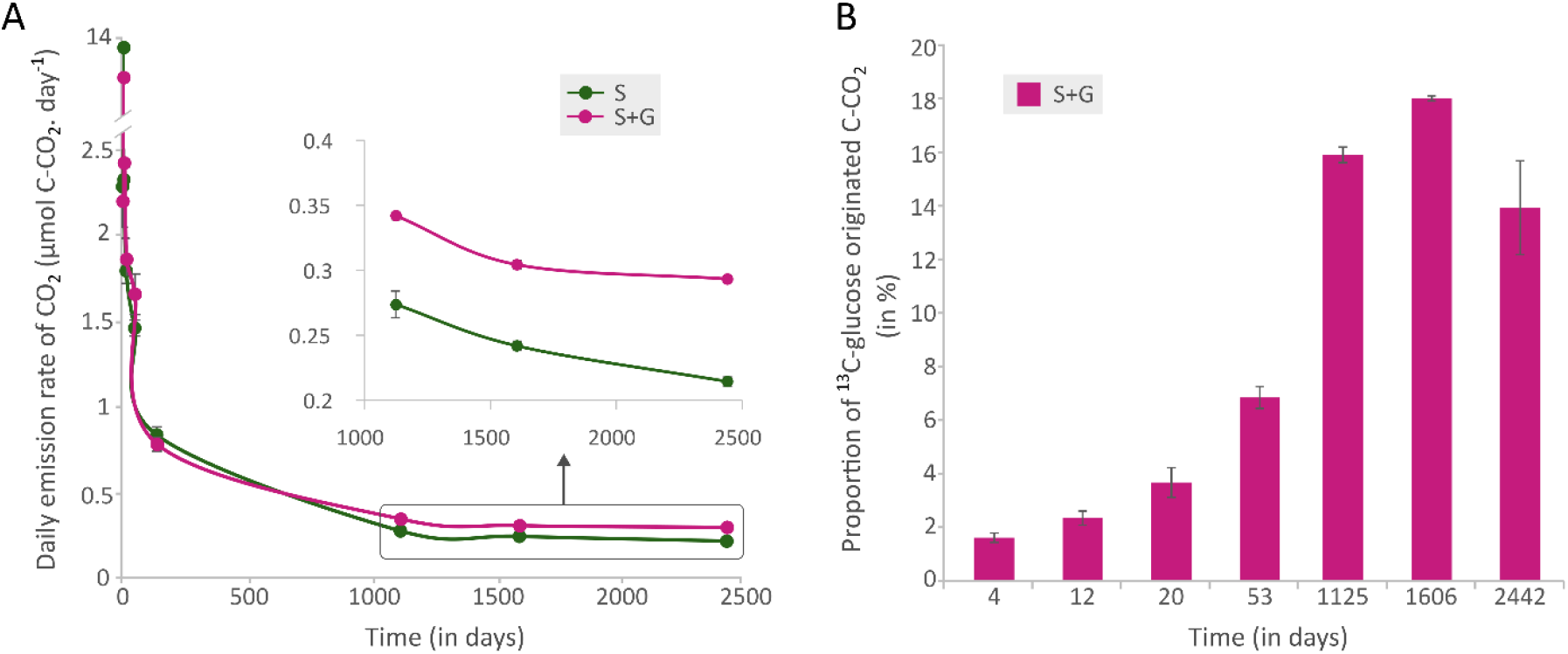
(A) Daily rate of total CO_2_ emissions and (B) percentage of glucose-derived C in daily CO_2_ emissions from irradiated soils with (S+G) and without (S) addition of ^13^C-labelled glucose in a long-term experiment (> 6 years). The inset in the upper right of panel A is an enlargement of the curves between 1125 and 2442 days. Standard deviations are shown in all plots.

Statistical modelling was used to analyse the kinetics of the CO_2_ emission rate and to test for the existence of different catalyst types and lifetimes supporting non-cellular catalytic activities (**Appendix S1: Section S6)**. The decrease in the CO_2_ emission rate over the first four days was excluded from the analysis in order to focus on the long-term behaviour of the catalytic activities. Six different models (**Figure 6**) were fitted to the CO_2_ emission rate of the glucose-amended soil to minimise the role of substrate limitation in the decrease in CO_2_ emission rate over time. Comparisons of these competing models using the Akaike’s information criterion led to the selection of two models (Models 5 and 6, **Figure 6**) that considered two different catalytic processes with contrasting persistence. For the catalytic process with the shortest persistence, the two models converge on a half-life of 71 days. This half-life is within the range of those observed for enzymes stabilized on soil particles such as clays (**Yan *et al*. 2010; Maire *et al*. 2013; Fanin *et al*. 2020**). For the second process, our results do not allow us to distinguish one model as more relevant than the other. Model 5 assumes that the catalytic process has a limited lifetime, with a half-life of seven years. Model 6 assumes that the catalytic process is completely stable over time. Given that enzymes, even when stabilised on soil particles, have a half-life of well under seven years (**Yan *et al*. 2010; Maire *et al*. 2013; Fanin *et al*. 2020)**, both models suggest that the most persistent non-cellular catalytic process cannot be mediated by enzymes.

**Figure 6:**
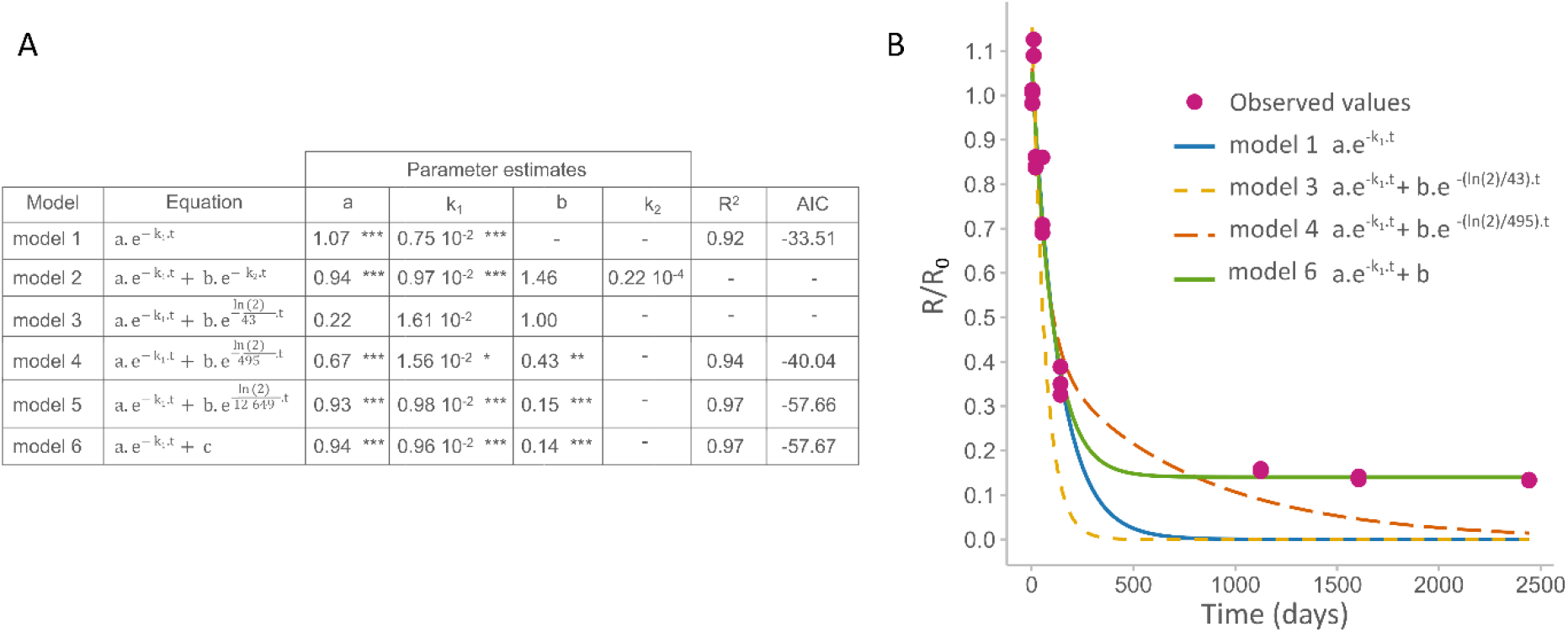
Modelling of the long-term dynamics of the daily rate of CO_2_ emission observed in irradiated soil supplemented with glucose (S+G). Model equations and statistical results obtained after fitting the models to the data. The models were fitted using the *nls* function in the lme4 R package. For models 2 to 5, the “port” algorithm was used as a nls parameter to constrain the estimates of k_1_ to a value greater than the decay rate of the second modelled activity kinetics. The absence of an asterisk after the estimate indicates a nonsignificant result. (**B**) Long-term dynamics of the daily CO_2_ emission rate. The release of CO_2_ over time (R) is expressed relative to the initial release of CO_2_ (R_0_). Each line represents the fit of each kinetic model to the measured daily CO_2_ emission rate data (pink circles).

We propose that persistent catalytic processes in sterilised soils involve the activity of highly enduring catalysts such as soil organominerals and/or minerals. Although this hypothesis may have been met with some scepticism a decade ago, recent studies convincingly demonstrate the previously overlooked but crucial role of reactive minerals in the processing and transformation of soil OM (*e*.*g*., **Wu *et al*. 2023**). These durable catalysts can facilitate OM transformation by multiple processes, including electrolytic and/or hydrolytic decomposition of macromolecules, heterogeneous oxidation, and direct oxidation mediated by transition elements (*e*.*g*., Fe or Mn) (**Kleber *et al*. 2021, Wu *et al*. 2023**). Redox-sensitive minerals, such as biotite, magnetite, Fe (II) sulfides, and Mn oxides can induce the hydrolytic decomposition of large macromolecular organic molecules into small molecules and can also efficiently catalyse the transformation of OM into new molecules (**Kleber *et al*. 2021**). For instance, in floodplain soils, reactive Fe (II) has been shown to be the main driver of OM oxidation, overtaking microbial activity (**Naughton *et al*. 2023**). In addition, some nano-sized mineral particles, called ‘nanozymes’, exhibit intrinsic enzyme-like activities, similar to those derived from biological sources, effectively catalysing the transformation of OM (**Wei *et al*. 2013**). For example, magnetite nanoparticles have been shown to have intrinsic enzyme-like activity that is similar to that of natural peroxidases, facilitating the oxidation of OM (**Gao *et al*. 2007**). Moreover, electron transfer, which we have demonstrated in sterile soils (**Figure 4**) and which occurs during mineral-mineral interactions in soil, can lead to the oxidation or reduction of OM compounds (**Qian *et al*. 2023**).

## CONCLUSION

The present study provides the first compelling evidence for EXOMETs, demonstrating that soil respiration can occur spontaneously without cellular compartmentalisation. In sterile soils, electron fluxes operate in conjunction with a series of time-ordered reactions, controlled by the nature of the substrate, that produce and consume exometabolites and lead to the complete oxidation of organic substrates with oxygen as the final electron acceptor. We show that different pools of catalysts are involved in these reactions (**Fig. 6**). In the short-term phases of our experiments (< months), EXOMETs are primarily induced by short-lived catalysts, likely cellular enzymes, which are continuously released by the turnover of organisms in natural soils. Soil sterilization, which triggers a peak in enzyme and substrate release and suppresses substrate uptake by living cells, enhances EXOMETs and has led to previous estimates of the potential contribution of EXOMETs to soil CO2 emissions of 12-50%, depending on soil properties (**Maire *et al*. 2013; Kéraval *et al*. 2018**). However, quantifying the contribution of EXOMETs to natural soil functioning remains a challenge that deserves further study. Collectively, these findings indicate that two types of interacting metabolisms, the respiration of living organisms and the EXOMETs, must be explicitly considered when studying biogeochemical cycles, as they are unlikely to obey the same laws and respond differently to environmental factors. The metabolism of living organisms is designed to meet the nutritional and reproductive needs of organisms that have limited lifespans, narrow physiological constraints (e.g. temperature, humidity), and are shaped by evolutionary processes. EXOMETs are not driven by ecological goals and are highly resistant to toxicity, high temperature, and pressure (**Maire *et al*. 2013; Blankinship *et al*. 2014**).

Our study also shows that non-living soils are unusual catalysts, stable and self-sustaining for periods much longer than the half-life of enzymes stabilised on soil particles (**Fig. 6**). In the latter phase of experiments (> months, years), EXOMETs depend on enduring non-enzymatic catalysts (e.g. minerals). It is striking that stochastic processes (*e*.*g*. random fluctuations in metabolite production and consumption rates, passive movements, or low exchange rates of metabolites) have a limited influence on the dynamics of metabolite assemblages in sterile soils, even in this last phase of experiments, showing that soil matrix shapes transformations with properties similar to biochemical transformations, given that deterministic processes (*e*.*g*. preferential consumption of thermodynamically more available compounds) are a feature of the latter **(Danczack *et al*. 2020**). This perspective offers interesting avenues for exploring fundamental questions of prebiotic chemistry, particularly with regard to the ability of minerals to catalyse sequences of metabolic reactions, which is the subject of debates. Some argue that, given the complexity of the metabolic pathways, it is unlikely that metabolism-like chemical reaction sequences could be catalysed by simple environmental catalysts (**Lazcano and Miller 1999; Anet 2004**). Others argue that the topology of glycolysis and the Krebs cycle is rooted in non-enzymatic chemistry, and that the precursors of these metabolic pathways were catalysed by abundant metal ions available on the primordial planet (**Keller *et al*. 2017, Ralser 2018**). For example, Fe^2+^ has been shown to be capable of catalysing 29 reactions involved in glycolysis and the pentose phosphate pathway in living organisms under conditions similar to those presumed in the Archaean ocean (**Keller *et al*. 2014**). Our study which shows non-enzymatic promotion of multiple reactions in successive sequences supports a scenario of non-enzymatic metabolism-like networks on the primitive Earth and their persistence in contemporary soil ecosystems.

## Supporting information

Supplemental materials for "Non-living respiration: another breath in the soil?" Bouquet et al. 2025

## ACKNOWLEDGEMENTS

Clémentin Bouquet was supported by PhD fellowship from the French Ministry of Education and Research. The authors would like to thanks the financial support of (i) the CNRS through the Mission pour les Initiatives transverses et interdisciplinaires (MITI) and through Projet Exploratoire Premier Soutien (PEPS) and the funding of the ISO-EXOMET and EXCEED projects, (ii) CAP 20-25 Isite of the University Clermont Auvergne and the funding of the EXOMET project.

## Authors Contribution

SF, GA, BK and ACL have designed the work. All authors contributed to the experimental data acquisition. SF, MT, GA, CB and ACL contributed to data set analysis and interpretation of data. CB and ACL drafted the work, and all authors have revised it. ACL and SF supervised the project.

