## Supplemental materials for "Non-living respiration: another breath in the soil?" Bouquet et al. 2025 for "Non-living respiration: another breath in the soil?"

### Appendix S.1

### Non-living respiration: another breath in the soil?

Clémentin Bouquet[1*], Benoit Kéraval[1*], Mounir Traikia[2,3], Gael Alvarez[4], Fanny Perrière[1], Anne-Hélène Le Jeune[1], Hermine Billard[1], Jonathan Colombet[1], Sandrine Revaillot[4], Sébastien Fontaine[4#], Anne-Catherine Lehours[1#⁑]

[1]Université Clermont Auvergne, CNRS, LMGE, F-63000 Clermont-Ferrand, France
[2]Université Clermont Auvergne, CNRS, ICCF, F-63000 Clermont-Ferrand, France
[3]Université Clermont Auvergne, PFEM, MetaboHUB Clermont, Clermont-Ferrand, France
[4]Université Clermont Auvergne, INRAE, VetAgro Sup, UMR Ecosystème Prairial, 63000 Clermont-Ferrand, France

* and # : These authors have contributed equally to this work

---

#### 1- Supplementary information: full material and methods (Section S1 to S6)

#### 2- Supplementary table

**Table S1: Comparisons of voltages recorded between and within different fuel cell conditions (soil+water (SW); soil+sodium pyruvate (SP), soil+sodium chloride (SC)) at different times of analysis and after the first and second injections of solutions. Comparisons were based on two-way repeated measures Anova with Sidak's post hoc test for multiple comparisons.**
**(A)** Comparison test between (SW) and (SP) at different times after the first injection, **(B)** Comparison test between (SW) and (SP) between different times after the second injection, **(C)** Comparison test at different times for (SW) after the first injection, **(D)** Comparison test at different times for (SW) after the second injection, **(E)** Comparison test at different times for (SP) after the first injection, **(F)** Comparison test at different times for (SP) after the second injection, **(G)** Comparison test between (SW) or (SP) and (SC) at different times.

#### 3- Supplementary figures

**Figure S.1: Soil sterility tests**
**(A)** Number of viable cells counted by flow cytometry (FCM) after live/dead staining either on soil extracts or after inoculation of culture medium (Luria Bertani (LB), malt extract (ME)) with 1 g of non-sterilised or sterilised bulk soil. The values and standard errors are shown on the graph for each condition **(B to D)** Examples of transmission electron microscopy photographs of non-sterilised (B), sterilised (C) and sterilised and $^{13}$C-glucose amended bulk soil (D). Green arrows in the photograph (D) indicate identifiable cell structures, the scale bars represent 1 µm (magnification x30000). **(E to G)** Examples of cytograms obtained for non-sterilised (E), sterilised (F), sterilised and $^{13}$C-glucose amended (G) soils after live-dead staining and FCM analysis. Further information on the sterility tests performed can be found in section S2 of the supplementary material. The propidium iodide fluorescence and the SybrGreen fluorescence are on the PE-Texas-Red-A and FITC-A axes, respectively.
T+: positive control, LB: Luria Bertani medium, ME: Malt extract medium, (S): sterilised soil microcosms, (S+G): sterilised soil amended with $^{13}$C-glucose microcosms, LS: non-sterilised soil, (G-IS): sterilised soil amended with $^{13}$C-glucose, (C-IS): sterilised soil amended with $^{13}$C-citrate.

**Figure S2: $^{1}$H nuclear magnetic resonance (NMR) spectra in the 0-10 ppm range for non-sterilised soil (LS) soil sterilised by *gamma irradiation* (IS) at different sampling dates**.
T1= 0.2 day, T2=3 days, T3=6 days, T4=17 days, T5=100 days, T6=163 days.

**Figure S3: Heat map showing relative bucket intensities of the water extractable fraction of exometabolites with molecular weight < 3 kDa at T1 (0.2 day) and T6 (163 days) in (A) irradiated soil (IS), (B) irradiated soil amended with $^{13}$C-citrate (C-IS) and in (C) irradiated soil amended with $^{13}$C-glucose (G-IS).**
Relative abundance is normalised to the concentration of the reference molecule during the $^{1}$H-NMR processing. Shades of red and blue indicate increasing and decreasing intensities, respectively. The colour scale represents the magnitude of change. Individual samples are shown on the horizontal axis and buckets on the vertical axis. Euclidean distance metric and Ward's clustering method were used for hierarchical clustering. Automatic normalisation of the bucket scale was also applied. The shades of red and blue indicate increasing and decreasing intensities, respectively

**Figure S4: Principal component analysis (PCA) of the temporal dynamics of the water-extractable fraction of exometabolites with molecular weight < 3 kDa in sterilised soil amended with (A) $^{13}$C-citrate and (B) $^{13}$C-Glucose.**
Each PCA point corresponds to one sacrificed replicate (4 replicates per sampling date). Coloured ellipses indicate 95 % confidence regions. The two-dimensional plot of the PCA shows that the first two principal components (PC) account for > 96.7 % of the variance explained with the first principal component (PC1) retaining > 78.4 % of the variance.
T1= 0.2 day, T2=3 days, T3=6 days, T4=17 days, T5=100 days, T6=163 days

**Figure S5: Principal component analysis (PCA) of the temporal dynamics of the water extractable fraction of exometabolites with molecular weight < 3 kDa in non-sterilised soil (LS) samples.**
Each point in the PCA corresponds to a sacrificed replicate (4 replicates per sampling date). Coloured ellipses indicate 95 % confidence regions. The two-dimensional PCA score plot shows that the first two principal components (PC) accpunt for 88.3 % of the variance explained, with the first principal component (PC1) retaining 69.5 % of the variance.
T1= 0.2 day, T2=3 days, T3=6 days, T4=17 days, T5=100 days, T6=163 days.

### 1. Supplementary information : full material and methods (Sections S1 to S6)

#### Section S1: Soil sampling and preparation of soil microcosms

Soil samples were taken from the 5-20 cm soil layer at the Theix site (Massif Central, France). The soil is a sandy loam Cambisol developed on granitic rocks and has the following characteristics: $pH_{water}$ = 6.2; carbon content = 39 gC. $kg^{-1}$; clay = 26 %; silt = 25 %; sand = 49 %; Cation Exchange Capacity (CEC) = 21.5 cmol. $Kg^{-1}$. For detailed information on the sampling site and on soil characteristics, see **Fontaine *et al.* (2007)** and **Kéraval *et al.* 2018.** Fresh soil samples were mixed, sieved at 2 mm and dried to 5 % (w/w of water per dry soil). Irradiated soil samples were sterilised by *gamma irradiation* at 45 kGy ($^{60}$Co, IONISOS, ISO14001, France). This sterilisation treatment effectively (i) suppresses active cells from soils (**Maire *et al.* 2013; Kéraval *et al*. 2016; McNamara *et al.* 2003**), (ii) limits the impact of sterilisation treatments on the physicochemical soil properties (**McNamara *et al.* 2003, Lees *et al.* 2018**), and (iii) yields similar respiration rates over a 15-day incubation period to soil sterilised by both *γ-irradiation* and autoclaving (**Kéraval *et al.* 2016**).

**S1A- Preparation of soil microcosms for the medium-term experiment (6 months):** The non-sterilised soil (LS) and irradiated soil (IS) microcosms consisted of 10 g sieved soil samples (dry mass) placed in 120 mL sterile glass vials capped with butyl rubber stoppers and sealed with aluminum crimps. Sterile solutions of uniformly labelled (*i.e.* all labelled positions) $^{13}$C-glucose (CAS: 110187-42-3, Cambridge Isotope Laboratories) and $^{13}$C-citrate (CAS: 287389-42-9, Aldrich) were prepared and either not amended (IS) or added (5 % $^{13}$C labelled) to the G-IS and C-IS microcosms, respectively, for a final concentration of glucose and citrate in a soil microcosm of 3 mgC. $g^{-1}$. All microcosms were flushed with sterilised free $CO_2$ gas (80 % $N_2$, 20 % $O_2$) moistened with sterilised water (LS and IS) or sterilized of $^{13}$C-glucose and $^{13}$C-citrate solutions (G-IS and C-IS) to a water content of 30 % and incubated in the dark at 20°C for 163 days. Four independent microcosm replicates per sampling date and treatment (IS, G-IS, C-IS) were incubated for the irradiated soils. Vials were sampled at T1 = 0.2 day, T2 = 3 days, T3 = 6 days, T4 = 17 days, T5 = 100 days and T6 = 163 days of incubation. All manipulations were carried out under sterile conditions and each sampling was destructive to avoid contamination of the soil microcosms.

**S1B- Preparation of soil microcosms for the long-term experiment (> 6 years):** Experimental microcosms consisted of 20 g of γ-irradiated soil incubated with 6 mL of water (S treatment) or 5 mL of water and 1 mL of $^{13}$C-labelled glucose solution (S+G treatment). The glucose solution contained 60 mg C-glucose $mL^{-1}$ and was prepared by mixing unlabelled glucose with $^{13}$C-labelled glucose ($C_6$ atom% $^{13}$C = 99%) to give a final $\delta^{13}$C of 3712 ‰. The glucose solution was sterilised by filtration at 0.2µm. Three replicates were made per treatment. The volume of solutions was adjusted to incubate soils at a water potential of -100 kPa. The soil microcosms were incubated at 25°C for 2442 days. All manipulations were performed under sterile conditions.

#### Section S2: Soil sterility tests

**S2A- Extraction of cells from soils:** At the end of the incubation period (t = 163 days and t= 2442 days for the medium-term and long-term experiment, respectively), the cells were separated from soil particles for all soil microcosms. One gram of soil was mixed with 10 mL pyrophosphate buffer (PBS 1X, 0.01 M $Na_4P_2O_7$) and shaken on ice at 70 rpm on a rotary shaker for 30 min. After shaking, the solution was sonicated 3 times (1 min each) in a water bath sonicator (Fisher Bioblock Scientific 88156,

320W, Illkirch, France). The largest particles were removed by centrifugation (800 × g, 1 min), and the supernatant was stored at 4°C for 2 h before quantification analysis.

**S2B- Live / dead cell staining coupled to flow cytometry analysis:** Samples were diluted in filtered Tris-EDTA (TE) buffer (<0,2 µm) and stained with 1X Sybr Green I (S7585, Invitrogen), a cell-permeable DNA dye that stains all live and dead cells. Cells were also stained with 10 µg/mL Propidium Iodide (P4864, MERCK KGaA), a cell-impermeable DNA dye that exclusively stains dead cells (**Crowley *et al.* 2016**), for 15 min in the dark at room temperature. Flow cytometry measurements were performed using a FACSAria Fusion SORP (BD Biosciences) equipped with a 70 µm nozzle and a 1.5 neutral density filter in a two lasers configuration depending on the dyes used (488 nm, 50 mW ; and 561 nm, 50 mW). The respective fluorescence emissions were collected with long pass (LP) and band pass (BP) filter sets : 502 nm LP / 530/30 nm BP (SG) ; 600nm LP / 610/20 nm BP (PI). The threshold was set at the minimum fluorescence on the SybrGreen I parameter. Data were acquired and analysed on logarithmic scales using FACSDiva 8 software (BD Biosciences).

**S2C- Cultural approaches:** For the long-term experiment (> 6 years), soil extracted at the end of the experiment (t= 2442 days) from the different (S) and (S+G) microcosm replicates were plated on Luria-Bertani (LB) and Malt extract (ME) agar plates to check whether colonies of prokaryotes or fungi formed on the surface of the agar plates. After one week of incubation at 25°C, no colonies were visible on the agar media.
In parallel, at t = 2442 days, 1 g of bulk soil samples from each of the (S) and (S+G) microcosms and for non-sterilised soil (used as positive control) were placed in flasks containing LB and ME broth. Cultures were incubated for 120 hours at 25°C with shaking. Cultures were analysed by flow cytometry after live/dead staining (**see section S2B**).

**S2D- Transmission electron microscopy (MET):** For the long-term experiment (> 6 years), at the end of the incubation period (t = 2442 days), bulk soil samples were analysed by TEM to check whether cells with preserved morphology were detectable in the soil aggregates, as previously described (**Kéraval *et al.* 2016**). Briefly, ultrathin soil sections (90 nm thick) were observed by TEM. Each step of the soil inclusion protocol was followed by centrifugation (12000 ×g, 2 min) to pellet the soil samples. Aliquots of the soil samples (0.05 g) were fixed for 1 h in 1.5 mL of a cacodylate buffer, pH 7.4 (0.2 M cacodylate, 6 % glutaraldehyde, and 0.15 % ruthenium red). The soil was washed three times with 0.1 M cacodylate buffer for 10 minutes. Post fixation was carried out with the 0.1 M cacodylate buffer containing 1 % osmic acid. To facilitate further penetration of propylene oxide, the soil was dehydrated through a gradient of ethanol: 50 % ethanol (3 × 5 min), 70 % ethanol (3 × 15 min) and 100 % ethanol (3 × 20 min) solutions. To improve resin permeation, the sample was incubated in a propylene oxide bath (3 × 20 min). To allow the sample to soak the resin, the soil sample was incubated overnight in a bath containing propylene oxide and Epon 812 resin (ratio 1 : 1) and secondary eliminated by flipping. After polymerisation of the cast resin on the soil specimens (48 h, 50 ◦C), the narrower parts of the moulded and impregnated aggregates were pyramided using a Reichert TM60 ultramill and finally, ultrathin sections (90 nm) were made using a diamond knife (Ultra 45◦, MF1845, DiATOME Ltd., Biel/Bienne, Switzerland; Ultramicrotome Ultracut S, Reichert Jung Leica, Austria). Soil sections were collected on 400-mesh Cu electron microscope grids supported by carbon-coated Formvar film (Pelanne Instruments, Toulouse, France). Each grid was negatively stained with uranyl acetate (2 %) for 30 s, rinsed twice with 0.02 µm distilled water, and dried on filter paper. Soil ultrathin sections were analysed using a JEM 1200-EX TEM (JEOL, Akishima, Japan).

**Section S3: Non-targeted metabolite profiling using proton nuclear magnetic resonance spectroscopy ($^1H$ NMR)**

**S3A- Extraction of exometabolites from soil:** Exometabolite extraction was performed using the method developed by **Jenkins *et al.* 2017** with minor modifications. Briefly, four grams of soil were extracted under sterile conditions in 24 mL of sterile MilliQ water at 4°C for 1 h whith shaking on an orbital shaker (Stuart SB3). The tubes were then centrifuged at 4 °C for 5 min at 3,200 × g. The supernatants were collected in fresh 50 mL conical tubes and centrifuged again. The supernatants were then filtered using Swinnex filtration units fitted with 25mm diameter GF/F (Whatman) previously grilled at 450°C for 4h and rinsed with 30mL of sterile milliQ water. These water extractable metabolites were frozen at -20 °C, lyophilised to dryness (Christ ALPHA 1-2 LDplus) and resuspended in 1.5 mL ultrapure water (MilliQ system). The samples were filtered on a centrifugal filter (Amicon ultra-4 3 kDa) previously rinsed using 4 mL sterile 0.1N NaOH and 4 mL sterile MilliQ water. After centrifugation for 45 min (7,500 g; 4°C), the < 3 kDa exometabolites were lyophilised again and stored at -80°C until analysis.

**S3B- Analysis by proton nuclear magnetic resonance spectroscopy ($^1H$ NMR):**

- Recipes for preparing $^1H$ NMR buffer:

-Buffer 1 (300 mM phosphate buffer solution, pH=7.28): 100 mL D2O (CAS: 7789-20-0, Cortecnet), 1.601 g $KH_2PO_4$ (CAS: 7778-77-0, Fluka analytical), 3.176 g de $K_2HPO_4$ (CAS: 7758-11-4, Fluka analytical), 80 mg reference molecule (TSP-d4, CAS: 24493-21-8, Sigma-Aldrich) and 40 mg $NaN_3$ (CAS 26628-22-8, Sigma-Aldrich).
-Buffer 2 (EDTA-d12 solution at 10 mM): 28.9 mg EDTA-d12 (304 g.mol$^{-1}$, ethylenediaminetetraacetic acid 98 % atom d-12, CD650P1, CORTECNET), 10 mL buffer 1.
-Buffer 3: 10 mL buffer 2, 5.5 mL buffer 1 and 31 mL $D_2O$, final pH=7.25.

- Sample preparation: Each tube of dry sample was filled with 600 µL of buffer 3, vortexed and transferred to a 5 mm diameter NMR tube (InnovaChem SAS, B-500-5-7).
- $^1H$ NMR spectra acquisition: $^1H$ NMR spectra were recorded at 298K on a Bruker AVANCE III 500 MHz spectrometer equipped with a Bruker 5 mm Prodigy inverse cryoprobe probe TXI ($^1H$/$^{13}C$/$^{15}N$) with a z-gradient coil probe (Bruker Biospin GmbH). A one dimensional $^1H$ NMR spectrum was acquired for all samples using a ZGPR sequence (with low power presaturation of the water frequency). 128 scans were collected with a 90° impulsion time of 9.94 µs at a power of 8.5W, a relaxation time of 5 s, an acquisition time of 3.3 s, a spectral window of 20 ppm and data points from 65 K filled from zero to 131 K prior to Fourier transformation with a line broadening of 0.3 Hz. All processing was performed using Bruker TopSpin 4.0.7. The NMR facility is ISO9001 certified.
- Analysis of $^1H$ NMR spectra: NMR spectra were imported and analysed using NMRProcFlow 1.4.20 software (https://nmrprocflow.org/). Spectra were aligned locally and manually by study group. Bucketting was performed using a peak-by-peak method with variable bucket sizes. The data matrix was normalised to the reference molecule bucket and a signal-to-noise-ratio threshold of 3 was chosen. Principal component analysis (PCA) was performed using Metaboanalyst 5.0 software (https://www.metaboanalyst.ca/; **Pang et *al.* 2022**). Four replicate blanks were analysed for each sampling date. A blank is defined as a sterilised water sample that has undergone all exometabolite extraction steps, sample preparation and $^1H$ NMR analysis. The order in which the samples were analysed was completely randomised.

For comparisons between treatments (IS, C-IS, G-IS), a baseline correction was applied and, to reduce the effect of glucose or citrate on the overall variance, we excluded, from the NMR spectrum, the regions [3.20-s 3.30 ppm]; [3.35-3.94 ppm]; [4.63-4.68 ppm]; [5.21-5.25 ppm] corresponding to α-glucose, β-glucose and the region [2.48-2.76 ppm] corresponding to citrate on the basis of the values commonly used for these molecules (**Fan 1995**). We acknowledge that these zones may contain other molecules of interest.

- Molecular richness and *alpha* diversity: The concept of diversity is primarily used to describe organisms, but ecological diversity indices have been shown to be useful descriptors of the diversity of organic molecules in soils (**Davenport *et al.* 2023**). For the water extractable fraction of exometabolites with molecular weights < 3 kDa, we calculated molecular richness using the sum of non-null buckets in each sample and the α-diversity using the Shannon-Weiner index (**Magurran 1988**), based on the relative abundance of buckets calculated from the sum of and intensities of peaks. *Alpha* diversity indices were calculated using the QIIME workflow (**Caporaso *et al.* 2010**) on Galaxy (https://usegalaxy.fr/).
- Data and statistical analyses: Statistical analyses were performed using GraphPad Prism version 8.0.1 (GraphPad Software, Inc). For principal component analysis, clustering, and heat maps were performed using Metabonalyst 5.0 software ((https://www.metaboanalyst.ca/).

#### Section S4: Gas flux measurements and isotopic composition of $CO_2$

**S4A- Concentrations of $CO_2$ and $O_2$**: For the medium-term experiment, gas phase samples were taken from the vials LS, IS, C-IS and G-IS at 0.2, 3, 6, 17, 100 and 163 days of incubation to measure $CO_2$ and $O_2$ concentrations. For the long-term experiment, the accumulation of $CO_2$ was measured after 0.5, 3.5, 10, 19, 53, 142, 1125, 1606 and 2442 days of incubation in the gas phase of (S) and (GS) microcosms. Analyses were carried out using a Chrompack 438 gas chromatograph (Packard Instruments, Downersgrove, IL, USA).

**S4B- Isotopic composition of $CO_2$:** Oxidation of $^{13}C$-labelled substrate was specifically quantified by monitoring the production of $^{13}C$-labelled $CO_2$. The amount and isotope composition ($\delta^{13}C$) of $CO_2$ were quantified using a cavity ring-down spectrometer analyser coupled to a small sample injection module (Picarro G2101-i analyzer coupled to the SSIM, Picarro Inc., Santa Clara, California, USA). A volume of 20 mL of gas was sampled by the analyser. The $CO_2$ concentration in the gas samples ranged from 300 to 2000 ppm $CO_2$ in accordance with the operating range of the analyser. The $CO_2$ concentration and $\delta^{13}C$ of the gas samples were measured at a frequency of 0.5 Hz for 10 min. The value provided by the analyser is the integrated value during these 10 min of measurement. A reference gas with a known concentration of $CO_2$ and $\delta^{13}C$ was injected between the samples. Gas samples were diluted with synthetic air ($O_2$, 20%; $N_2$, 80%) when the $CO_2$ concentration exceeded the operating range of the analyser. More specifically for the long-term experiment, for each incubation period, the cumulative amount of $CO_2$ was divided by the duration of the period (in days) to calculate the average daily $CO_2$ emission rate. The daily $CO_2$ emission rate was expressed as C-$CO_2$ (µmole of C released as the form of $CO_2$) per day per incubation flask. After gas analysis, the flasks were flushed with synthetic air ($O_2$, 20%; $N_2$, 80%) moistened by bubbling water sterilised by filtration at 0.2 µm.

#### Section S5: Principle, design and fuel cell measurements

The principle of the fuel cell is based on a column of soil, the surface part of which is exposed to oxygen from the air, while the lower part is confined by the plastic tube (**Figure 4**). This design creates a decreasing oxygen gradient from the top of the cell (the cathode) to the bottom of the cell (the anode) similar to that found in soil. The oxygen gradient promotes the capture of the electrons produced from the oxidation of organic molecules by the anode and their transfer to the cathode. The protons released by the oxidation of organic molecules diffuse through the soil column to reach the cathode where the oxygen is reduced in water. The current induced by the transfer of electrons from the degradation of organic C to oxygen is measured using an amperemeter and a voltmeter. Soil contains a variety of organic and inorganic molecules that can respond to this oxygen gradient and generate electron flow by ways other than the oxidation of organic molecule (*e.g.* oxidation of $Fe^{2+}$ to $Fe^{3+}$). To demonstrate the link between the oxidation of organic molecule and electron flow, the current induced by the cell was measured with and without the addition of pyruvate.

**S5A- Soil matrix preparation:** Samples were collected from the 0-20 cm soil layer at the site of Theix (see **section S1**). Fresh soil samples were mixed, sieved at 2 mm, dried to 5 % (w/w of water per dry soil) and irradiated with gamma rays at 45 kGy ($^{60}Co$, IONISOS, ISO14001, France). The soils were packaged in small hermetically sealed plastic bags, which were sealed in other plastic bags (3 layers). The soils were stored at 20°C for 4.5 years before being used to construct the air breathing cells. This is a period long enough for cell debris to become invisible under electron microscopes (**Kéraval *et al.* 2016**), and for enzymes to be inactivated (**Maire *et al.* 2013**).

**S5B- Preparation of the fuel cell:** The fuel cell was constructed using a plastic tube (polypropylene, Falcon®) with a diameter of 3 cm, a height of 11 cm high and a capacity of 50 ml (**Figure 4**). The bottom of the tube and the cap were pierced, and a 2 mm diameter graphite rod was glued in place. These rods respectively collect and transfer electrons to the anode and cathode of the fuel cell. These rods were covered with carbon cloth (10 $cm^2$) to improve the contact surface with the soil reactor and electron collection/distribution. The tube with the bonded electrodes as well as the pieces of carbon cloth were autoclaved (121°C, 20 min) before the fuel cell was assembled.

**S5C- Assembly of the fuel cell:** All the manipulations were performed under a laminar flow hood. Three treatments were set up, including (1) sterilised soil with a sterile solution of sodium pyruvate (50 mM, Sigma Aldrich Reagent Plus ≥99.9% CAS: 113-24-6), (2) sterilised soil with sterile water and (3) sterilised soil with a sterile solution of sodium chloride (50 mM, R.P. Normapur analytical reagent ≥99.5 %). Treatments (2) and (3) were controls for treatment (1), which was designed to test the effect of a supply of readily degradable carbon on electron flow. Three replicates of each treatment were made. A volume of 16 mL of one of the three solution was poured into the fuel cell, then the 37.5 g of sterilised soil (fresh weight basis, 4% moisture) was poured in slowly to avoid air bubbles. The carbon tissue was placed on the surface of the wet soil. The tube cover was screwed on and the cathode carbon electrode was threaded through the hole in the cover and placed on the soil surface.

**S5D- Electrochemical characterization of fuel cell:** All fuel cells were tested using a precision digital multimeter (UNI-T-DT830B) to measure voltage (V) and current (I). The open circuit potential, current and electrical power of the fuel cells connected to a 476 ohm resistor were measured at 0.08, 0.7, 1, 1.75, 3, 6, 9, 13.5, 19.5 days of incubation. Electromotive force and fuel cell resistance were also determined by connecting the fuel cell to decreasing resistances (from 330 to 11900 ohms) on day 9.

**S5E- Data and statistical analyses**: Analyses were performed with GraphPad Prism software version 8.0.1. Comparisons were based on two-way repeated measures Anova with Sidak's post hoc test for multiple comparisons.

#### <u>Section S.6:</u> Modelling the long-term dynamics of the daily rate of $CO_2$ emission observed in irradiated soil supplied with glucose (S+G)

The long-term dynamics of the $CO_2$ emission rate was analysed using statistical modelling. Six different models were fitted successively to the decrease in $CO_2$ activity over time to investigate whether the fast and slow phases of the decrease were due to activities of combined non-cellular catalysts with different kinetics.

The models were fitted to the $CO_2$ emission rate observed in the glucose-supplied mesocosms, where substrate-limiting conditions were minimised. In this context, the models were mainly designed to test the possible involvement of catalysts of different types and lifetimes responsible for OM oxidation in sterilised soils.

Model 1 describes a simple exponential decrease of $CO_2$ activity with a constant decay rate k. Models 2 to 5 combine two exponentially decreasing $CO_2$ activities with fast ($k_1$) and slow ($k_2$) decay rates respectively. While the decay rate of the slow kinetics ($k_2$) was fitted to the data in model 2, in models 3 and 4 it was forced to values giving half-lives of 43 and 495 days corresponding to estimates of **Maire et al. (2013)** for long term stabilized enzyme pools. Similarly, the value of $k_2$ was forced in model 5 to a hypothetically extremely high half-life of 7 years never observed for an enzyme pool. Finally, model 6 combines a rapid exponential decrease in $CO_2$ emission rate with a constant slow activity maintained throughout the long-term dynamics.

The models were fitted using the *nls* function from the lme4 R package (https://CRAN.R-project.org/package=lme4). For models 2 to 5, the "port" algorithm was used as a parameter of *nls* to constrain estimates of $k_1$ to a value higher than the decay rate of the second activity kinetics modelled. The **Figure 6A** shows the models equations and all the statistical results obtained after fitting the models to the data. Only models 1, 4, 5 and 6 produced significant estimates. The likelihood of these competing models was then compared using the Akaike information criterion (AIC) in order to isolate the best models. **Figure 6B** shows the observed and simulated data by models 1, 3, 4 and 6.

### 2. SUPPLEMENTARY TABLE

**Table S1: Comparisons of voltages recorded between and within different fuel cell conditions (soil+water (SW); soil+sodium pyruvate (SP), soil+sodium chloride (SC)) at different times of analysis and after the first and second injections of solutions. Comparisons were based on two-way repeated measures Anova with Sidak's post hoc test for multiple comparisons.**

**(A)** Comparison test between (SW) and (SP) at different times after the first injection, **(B)** Comparison test between (SW) and (SP) between different times after the second injection, **(C)** Comparison test at different times for (SW) after the first injection, **(D)** Comparison test at different times for (SW) after the second injection, **(E)** Comparison test at different times for (SP) after the first injection, **(F)** Comparison test at different times for (SP) after the second injection, **(G)** Comparison test between (SW) or (SP) and (SC) at different times.

#### (A) (SW) versus (SP) after the first injection

| N° | Time after injection (days) | Mean diff. | 95.00% CI diff. | Summary | P. value |
|---|---|---|---|---|---|
| T1 | 0,083333333 | -26,09 | -32,58 to -19,61 | **** | <0,0001 |
| T2 | 0,6875 | -25,8 | -32,28 to -19,32 | **** | <0,0001 |
| T3 | 0,97916 | -28,64 | -35,12 to -22,16 | **** | <0,0001 |
| T4 | 1,75 | -31,45 | -37,93 to -24,97 | **** | <0,0001 |
| T5 | 2,9583333 | -32,02 | -38,50 to -25,54 | **** | <0,0001 |
| T6 | 5,9270833 | -17,39 | -23,87 to -10,91 | **** | <0,0001 |
| T7 | 8,916666 | -7,837 | -14,32 to -1,356 | ** | 0,0091 |
| T8 | 13,61249933 | -4,577 | -11,06 to 1,904 | ns | 0,3543 |
| T9 | 19,57083267 | -2,907 | -9,387 to 3,574 | ns | 0,8806 |
| T10 | 27,779166 | -1,56 | -8,040 to 4,920 | ns | 0,9986 |

#### (B) (SW) versus (SP) after the second injection

| N° | Time after first injection (days) | Time after second injection (days) | Mean diff. | 95.00% CI diff. | Summary | P. value |
|---|---|---|---|---|---|---|
| T11 | 77 | | -0,06667 | -2,224 to 2,091 | ns | >0,9999 |
| T12 | 77,01375 | 0,01375 | -2,633 | -4,791 to -0,4761 | * | 0,0101 |
| T13 | 77,0275 | 0,0275 | -2,667 | -4,824 to -0,5094 | ** | 0,009 |
| T14 | 77,05916667 | 0,059166667 | -3,733 | -5,891 to -1,576 | *** | 0,0002 |
| T15 | 77,10083333 | 0,100833333 | -4,167 | -6,324 to -2,009 | **** | <0,0001 |
| T16 | 78,05916667 | 1,059166667 | -8,733 | -10,89 to -6,576 | **** | <0,0001 |
| T17 | 79,93416667 | 2,934166667 | -10,83 | -12,99 to -8,676 | **** | <0,0001 |

#### (C) (SW) at different times after the first injection

| N° | Mean diff. | 95.00% CI diff. | Summary | P. value |
|---|---|---|---|---|
| T1 vs. T2 | 0,9943 | -4,767 to 6,755 | ns | 0,9998 |
| T1 vs. T3 | 0,6329 | -5,128 to 6,394 | ns | >0,9999 |
| T1 vs. T4 | -0,4125 | -6,174 to 5,349 | ns | >0,9999 |
| T1 vs. T5 | -1,379 | -7,140 to 4,382 | ns | 0,9975 |
| T1 vs. T6 | -3,236 | -8,997 to 2,526 | ns | 0,6398 |
| T1 vs. T7 | -1,882 | -7,643 to 3,879 | ns | 0,9772 |
| T1 vs. T8 | 0,2145 | -5,547 to 5,976 | ns | >0,9999 |
| T1 vs. T9 | 0,9111 | -4,850 to 6,672 | ns | >0,9999 |
| T1 vs. T10 | 1,564 | -4,197 to 7,326 | ns | 0,9936 |

#### (D) (SW) at different times after the second injection

| N° | Mean diff. | 95.00% CI diff. | Summary | P. value |
|---|---|---|---|---|
| T11 vs. T12 | 0,2333 | -1,568 to 2,035 | ns | 0,9994 |
| T11 vs. T13 | 0,2 | -1,601 to 2,001 | ns | 0,9998 |
| T11 vs. T14 | 0,2 | -1,601 to 2,001 | ns | 0,9998 |
| T11 vs. T15 | 0,2167 | -1,585 to 2,018 | ns | 0,9996 |
| T11 vs. T16 | 0,2 | -1,601 to 2,001 | ns | 0,9998 |
| T11 vs. T17 | 0,4 | -1,401 to 2,201 | ns | 0,9893 |

#### (E) (SP) at different times after the first injection

| N° | Mean diff. | 95.00% CI diff. | Summary | P. value |
|---|---|---|---|---|
| T1 vs. T2 | 1,291 | -4,471 to 7,052 | ns | 0,9985 |
| T1 vs. T3 | -1,916 | -7,677 to 3,846 | ns | 0,9744 |
| T1 vs. T4 | -5,763 | -11,52 to -0,002108 | * | 0,0499 |
| T1 vs. T5 | -7,301 | -13,06 to -1,540 | ** | 0,0059 |
| T1 vs. T6 | 5,466 | -0,2951 to 11,23 | ns | 0,0728 |
| T1 vs. T7 | 16,38 | 10,61 to 22,14 | **** | <0,0001 |
| T1 vs. T8 | 21,73 | 15,97 to 27,49 | **** | <0,0001 |
| T1 vs. T9 | 24,1 | 18,34 to 29,86 | **** | <0,0001 |
| T1 vs. T10 | 26,1 | 20,34 to 31,86 | **** | <0,0001 |

#### (F) (SP) at different times after the second injection

| N° | Mean diff. | 95.00% CI diff. | Summary | P. value |
|---|---|---|---|---|
| T11 vs. T12 | -2,333 | -4,135 to -0,5319 | ** | 0,0065 |
| T11 vs. T13 | -2,4 | -4,201 to -0,5986 | ** | 0,005 |
| T11 vs. T14 | -3,467 | -5,268 to -1,665 | **** | <0,0001 |
| T11 vs. T15 | -3,883 | -5,685 to -2,082 | **** | <0,0001 |
| T11 vs. T16 | -8,467 | -10,27 to -6,665 | **** | <0,0001 |
| T11 vs. T17 | -10,37 | -12,17 to -8,565 | **** | <0,0001 |

#### (G) (SP) or (SW) versus (SC) at different times

| N° | Time (days) | Comparison | Mean diff. | 95.00% CI diff. | Summary | P. value |
|---|---|---|---|---|---|---|
| T1 | 0,08 | SW vs. SC | -1,512 | -8,095 to 5,071 | ns | 0,9181 |
| | | SP vs. SC | 24,58 | 18,00 to 31,17 | **** | <0,0001 |
| T3 | 0,98 | SW vs. SC | -4,195 | -10,78 to 2,388 | ns | 0,3129 |
| | | SP vs. SC | 24,45 | 17,87 to 31,03 | **** | <0,0001 |
| T4 | 1,75 | SW vs. SC | -3,673 | -10,26 to 2,910 | ns | 0,4253 |
| | | SP vs. SC | 27,77 | 21,19 to 34,36 | **** | <0,0001 |
| T5 | 2,96 | SW vs. SC | -3,367 | -9,950 to 3,216 | ns | 0,4992 |
| | | SP vs. SC | 28,65 | 22,07 to 35,23 | **** | <0,0001 |
| T7 | 8,92 | SW vs. SC | -3,58 | -10,16 to 3,003 | ns | 0,4472 |
| | | SP vs. SC | 4,257 | -2,326 to 10,84 | ns | 0,301 |

### 3. SUPPLEMENTARY FIGURES

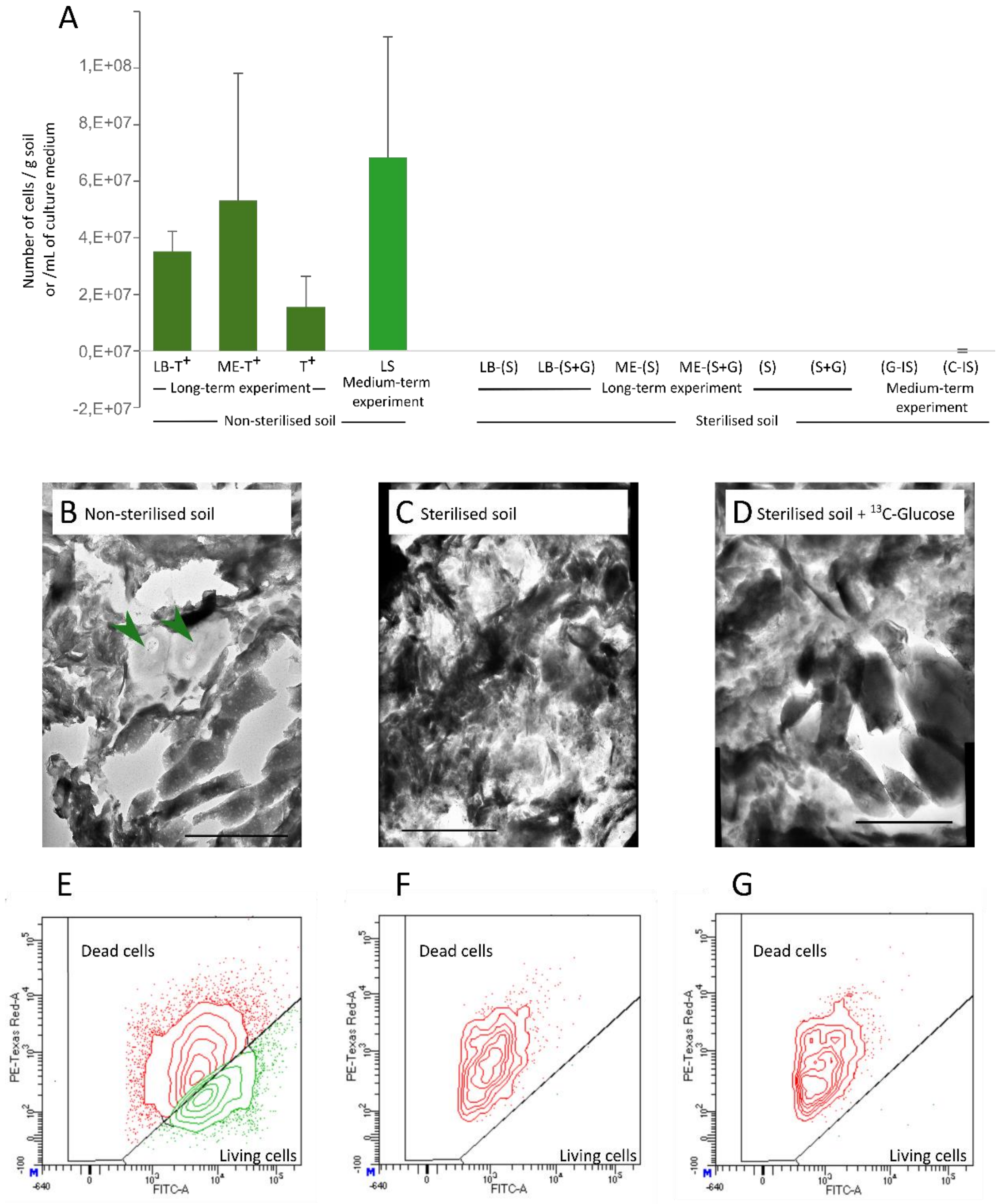


**Figure S.1: Soil sterility tests**

**(B)** Number of viable cells counted by flow cytometry (FCM) after live/dead staining either on soil extracts or after inoculation of culture medium (Luria Bertani (LB), malt extract (ME)) with 1 g of non-sterilised or sterilised bulk soil. The values and standard errors are shown on the graph for each condition **(B to D)** Examples of transmission electron microscopy photographs of non-sterilised (B), sterilised (C) and sterilised and $^{13}$C-glucose amended bulk soil (D). Green arrows in the photograph (D) indicate identifiable cell structures, the scale bars represent 1 µm (magnification x30000). **(E to G)** Examples of cytograms obtained for non-sterilised (E), sterilised (F), sterilised and $^{13}$C-glucose amended (G) soils after live-dead staining and FCM analysis. Further information on the sterility tests performed can be found in section S2 of the supplementary material. The propidium iodide fluorescence and the SybrGreen fluorescence are on the PE-Texas-Red-A and FITC-A axes, respectively.

T+: positive control, LB: Luria Bertani medium, ME: Malt extract medium, (S): sterilised soil microcosms, (S+G): sterilised soil amended with $^{13}$C-glucose microcosms, LS: non-sterilised soil, (G-IS): sterilised soil amended with $^{13}$C-glucose, (C-IS): sterilised soil amended with $^{13}$C-citrate.

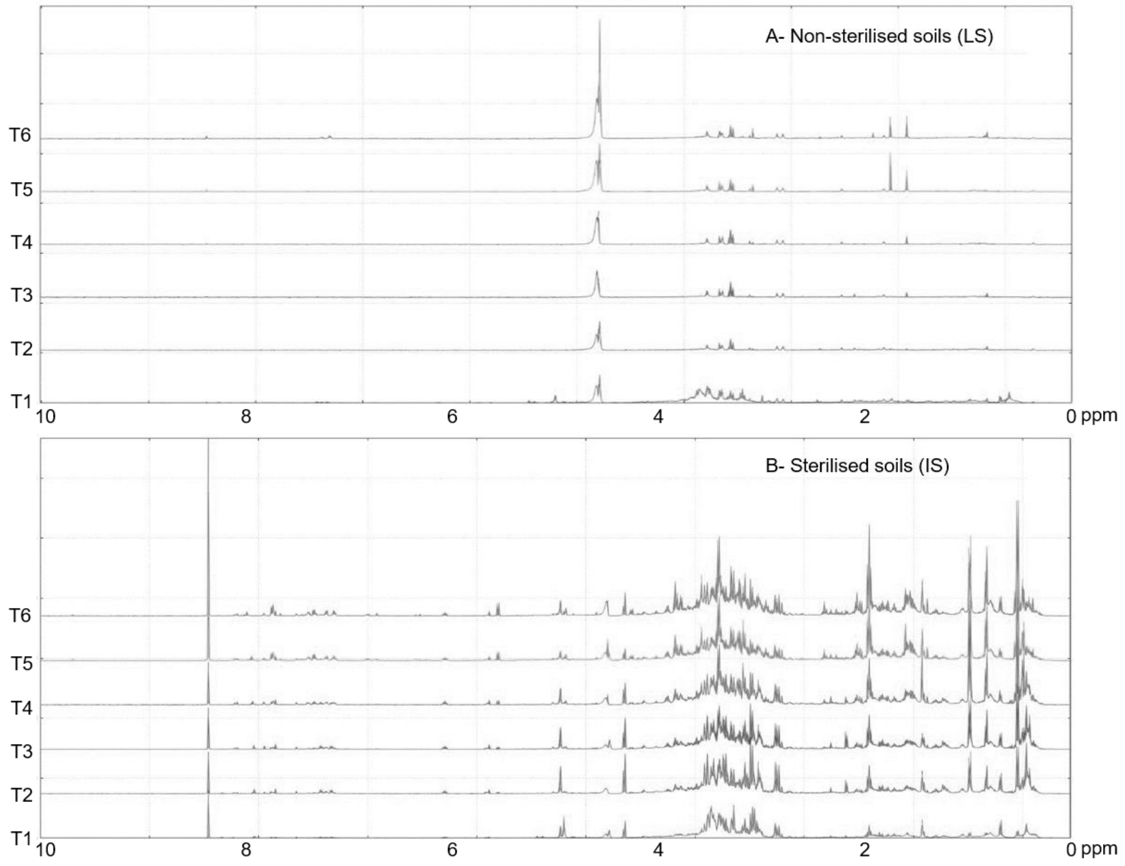


**Figure S2: $^{1}$H nuclear magnetic resonance (NMR) spectra in the 0-10 ppm range for non-sterilised soil (LS) soil sterilised by *gamma irradiation* (IS) at different sampling dates**.
T1= 0.2 day, T2=3 days, T3=6 days, T4=17 days, T5=100 days, T6=163 days.

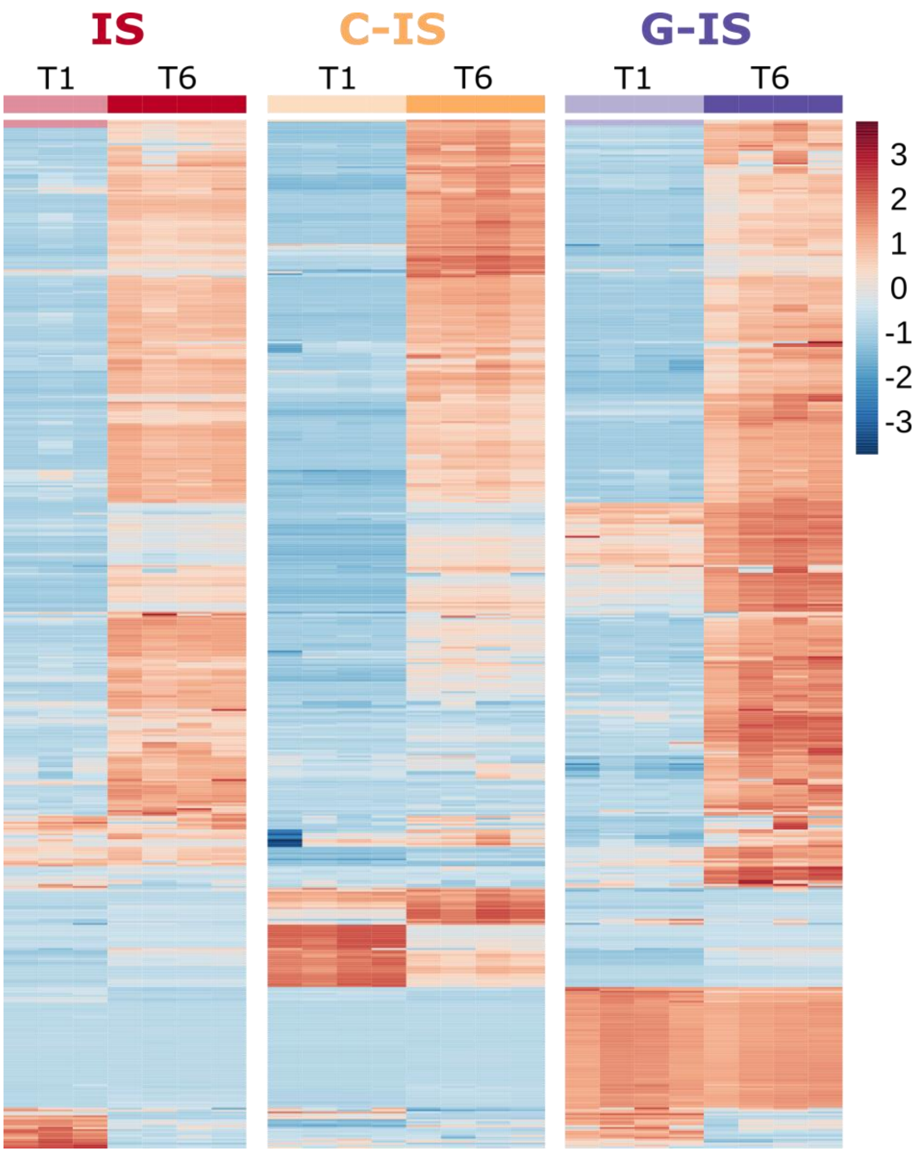


**Figure S3: Heat map showing relative bucket intensities of the water extractable fraction of exometabolites with molecular weight < 3 kDa at T1 (0.2 day) and T6 (163 days) in (A) irradiated soil (IS), (B) irradiated soil amended with $^{13}$C-citrate (C-IS) and in (C) irradiated soil amended with $^{13}$C-glucose (G-IS).**
Relative abundance is normalised to the concentration of the reference molecule during the $^{1}$H-NMR processing. Shades of red and blue indicate increasing and decreasing intensities, respectively. The colour scale represents the magnitude of change. Individual samples are shown on the horizontal axis and buckets on the vertical axis. Euclidean distance metric and Ward's clustering method were used for hierarchical clustering. Automatic normalisation of the bucket scale was also applied. The shades of red and blue indicate increasing and decreasing intensities, respectively

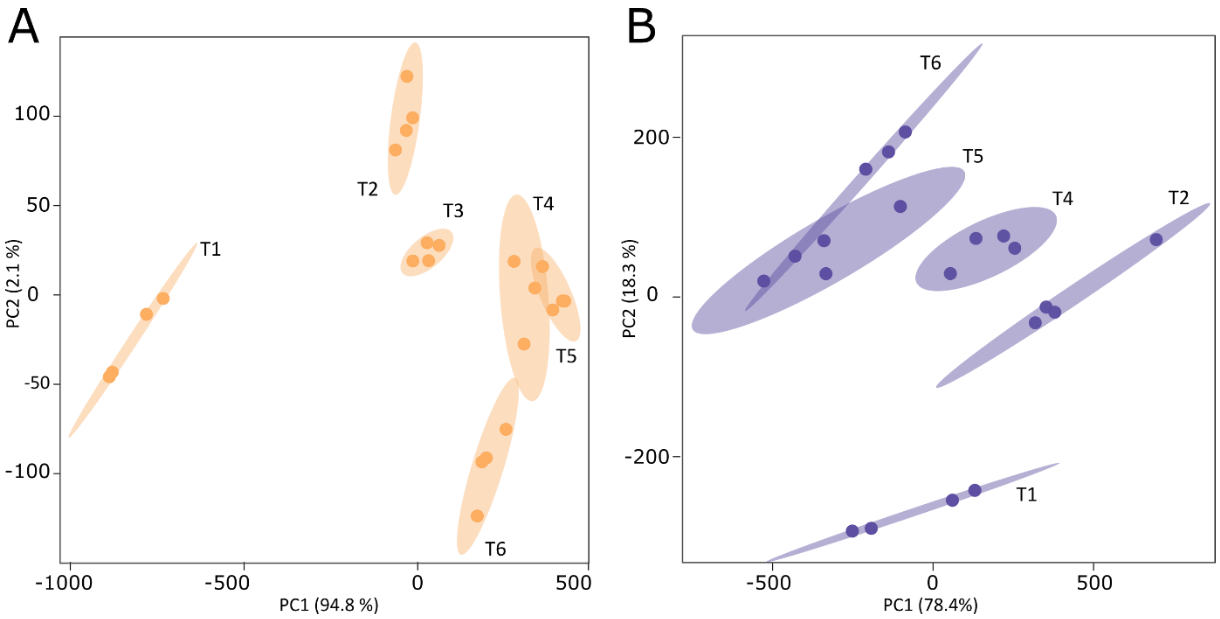


**Figure S4: Principal component analysis (PCA) of the temporal dynamics of the water-extractable fraction of exometabolites with molecular weight < 3 kDa in sterilised soil amended with (A) $^{13}$C-citrate and (B) $^{13}$C-Glucose.**

Each PCA point corresponds to one sacrificed replicate (4 replicates per sampling date). Coloured ellipses indicate 95 % confidence regions. The two-dimensional plot of the PCA shows that the first two principal components (PC) account for > 96.7 % of the variance explained with the first principal component (PC1) retaining > 78.4 % of the variance.

T1= 0.2 day, T2=3 days, T3=6 days, T4=17 days, T5=100 days, T6=163 days

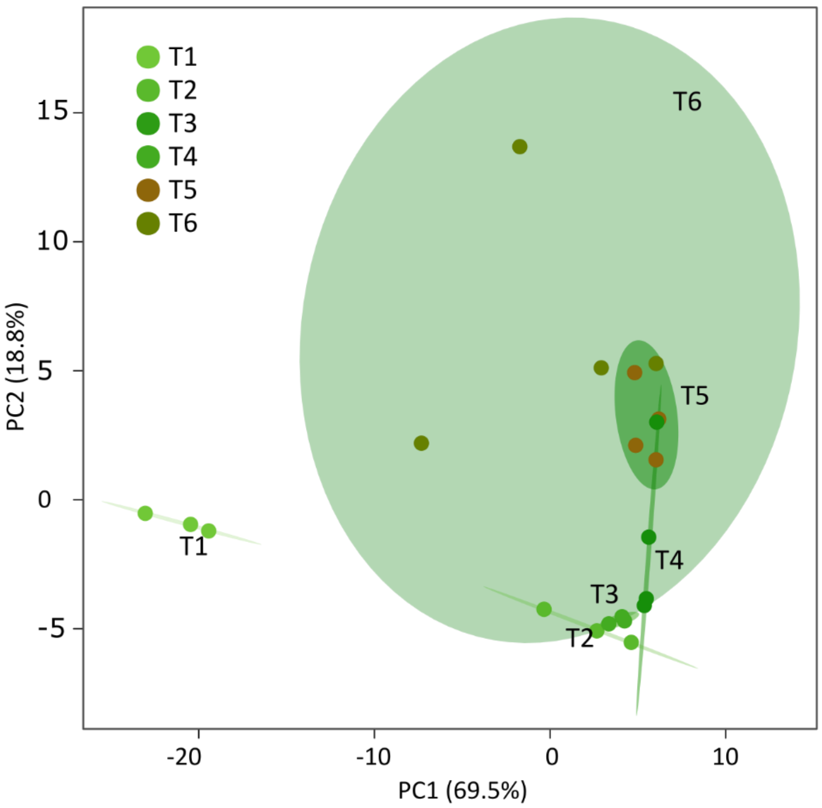


**Figure S5: Principal component analysis (PCA) of the temporal dynamics of the water extractable fraction of exometabolites with molecular weight < 3 kDa in non-sterilised soil (LS) samples.**
Each point in the PCA corresponds to a sacrificed replicate (4 replicates per sampling date). Coloured ellipses indicate 95 % confidence regions. The two-dimensional PCA score plot shows that the first two principal components (PC) accpunt for 88.3 % of the variance explained, with the first principal component (PC1) retaining 69.5 % of the variance.
T1= 0.2 day, T2=3 days, T3=6 days, T4=17 days, T5=100 days, T6=163 days.
